# Predicting expression divergence and its evolutionary parameters between single-copy genes in two species

**DOI:** 10.1101/2022.07.13.499803

**Authors:** Antara Anika Piya, Michael DeGiorgio, Raquel Assis

## Abstract

Predicting gene expression divergence and its evolutionary parameters is integral to understanding the emergence of new gene functions and associated traits. Whereas several sophisticated methods have been developed for these tasks, their applications are either limited to duplicate genes or require expression data from more than two species. Thus, here we present PiXi, the first machine learning framework for predicting expression divergence and its evolutionary parameters between single-copy genes in two species. In particular, PiXi models gene expression evolution as an Ornstein-Uhlenbeck process, and overlays this model with multi-layer neural network, random forest, and support vector machine architectures for making predictions. We show that PiXi has high power and accuracy in predicting gene expression divergence and its underlying parameters across a wide range of evolutionary scenarios, with the globally best performance achieved by a multi-layer neural network. Moreover, application of our best performing PiXi predictor to empirical data from single-copy genes residing at different loci in two species of *Drosophila* reveals that expression divergence occurs in approximately 20% of these positionally relocated genes and is driven by a combination of neutral and selective forces. Further analysis shows that several of these genes are involved in the electron transport chain of the mitochondrial membrane, suggesting that new chromatin environments may impact energy production in *Drosophila*. Thus, by providing a toolkit for predicting expression divergence and its evolutionary parameters between single-copy genes in two species, PiXi can shed light on the origins of novel phenotypes across diverse biological processes and study systems.

## Introduction

Determining whether and how gene functions have diverged between species is a problem of central importance in evolutionary genomics. In particular, researchers are often interested in assaying inter-species functional divergence for a specific set of genes, such as those that have undergone a mutation event or are involved in a biological process that is being studied [Gu, 1999, Lynch and Force, 2000, Gu, 2001, Kondrashov et al., 2002, Blanc and Wolfe, 2004, Li et al., 2005, Chain et al., 2008, Lopez-Bigas et al., 2008, Lynch and Wagner, 2008, Assis et al., 2012, Assis and Bachtrog, 2013, 2015, Assis, 2016, Fuller et al., 2016, Wheeler et al., 2016, Hart et al., 2018, Assis, 2019a, Jiang and Assis, 2019, Meng et al., 2019, Assis, 2021, Zhong et al., 2021, Jiang and Assis, 2022]. In these scenarios, the first major question to address is whether the functions of these genes are conserved or have diverged as a result of the mutation event or biological process under consideration. Second, one may want to know what evolutionary forces are responsible for functional conservation or divergence of these genes. Answering these questions is critical not only for learning about the functional divergence of a specific set of genes, but also for generating testable hypotheses about their contributions to the origins of complex phenotypes and species.

The classical approach to this common problem in evolutionary genomics is to quantify sequence divergence between orthologous genes, or those that arose from the same common ancestor, in related species [Gu, 1999, 2001, Kondrashov et al., 2002, Chain et al., 2008, Lopez-Bigas et al., 2008, Wheeler et al., 2016, Hart et al., 2018, Assis, 2019a, Zhong et al., 2021, Jiang and Assis, 2022]. Though such analyses enable estimations of the types and strengths of natural selection acting on a set of genes, they are limited in their ability to detect their functional divergence. Specifically, natural selection acts directly on gene functions, and therefore indirectly on their underlying sequences. With this in mind, several modern studies have assayed functional divergence from gene expression data [Blanc and Wolfe, 2004, Li et al., 2005, Chain et al., 2008, Assis et al., 2012, Assis and Bachtrog, 2013, 2015, Assis, 2016, Fuller et al., 2016, Perry and Assis, 2016, Hart et al., 2018, Assis, 2019a, Jiang and Assis, 2019, Meng et al., 2019, Zhong et al., 2021, Jiang and Assis, 2022], which are now widely available for many conditions (e.g., tissues, developmental stages, or disease states) in diverse species [Kapushesky et al., 2010, Consortium, 2012, Petryszak et al., 2013]. Because expression data can provide information about activity levels of a gene across multiple conditions, it is often considered an ideal proxy for function [Wray et al., 2003, Carroll, 2005, Nehrt et al., 2011, Assis and Bachtrog, 2013, De Smet et al., 2017]. Further, gene expression is easily quantified and compared, and also strongly correlated with a number of other important genic properties, including protein-coding sequence divergence [Makova and Li, 2003, Nuzhdin et al., 2004, Lemos et al., 2005, Hunt et al., 2012, Assis, 2014, Assis and Kondrashov, 2014, Mähler et al., 2017, Assis, 2019b] and protein-protein interactions [Bhardwaj and Lu, 2005, Lemos et al., 2005, Assis and Bachtrog, 2013, Assis and Kondrashov, 2014, Musungu et al., 2016, Mähler et al., 2017, Assis, 2019b].

In recent years, Ornstein-Uhlenbeck (OU) processes have been used to develop many sophisticated methods for modeling expression evolution of orthologous genes along phylogenetic trees [Hansen, 1997, Butler and King, 2004, Kalinka et al., 2010, Brawand et al., 2011, Perry et al., 2012, Rohlfs et al., 2014, Rohlfs and Nielsen, 2015, DeGiorgio and Assis, 2021]. Because OU processes model Brownian motion with a pull toward an optimal state, they have a natural application to evolution, in which phenotypic drift is analogous to Brownian motion, selection to pull, and the fittest phenotype to optimal state [Hansen, 1997, Butler and King, 2004]. Whereas most of these OU-based methods can also be used to assay expression divergence and its evolutionary parameters [Hansen, 1997, Butler and King, 2004, Brawand et al., 2011, Rohlfs et al., 2014, Rohlfs and Nielsen, 2015, DeGiorgio and Assis, 2021], they are limited in their applicability to problems generally encountered in evolutionary genomics. Specifically, these methods either require gene expression data from more than two species [Hansen, 1997, Butler and King, 2004, Brawand et al., 2011, Rohlfs et al., 2014, Rohlfs and Nielsen, 2015], which researchers typically do not have access to, or are tailored to genes that underwent duplication events [DeGiorgio and Assis, 2021]. Thus, there are currently few options for predicting expression divergence and its evolutionary forces between single-copy genes in two species.

Here we present PredIcting eXpression dIvergence (PiXi), an OU model-based machine learning framework for predicting expression divergence and its underlying evolutionary parameters between single-copy genes in two species. As in a recent method designed for duplicate genes, CLOUD [DeGiorgio and Assis, 2021], we choose machine learning for prediction due to several advantages over traditional likelihood ratio tests previously used for single-copy genes [Kalinka et al., 2010, Brawand et al., 2011, Perry et al., 2012, Rohlfs et al., 2014, Rohlfs and Nielsen, 2015]. For one, machine learning algorithms do not require assumptions to be made about evolution, instead discriminating among models based solely on features extracted from data [Hastie et al., 2009]. Second, training of machine learning algorithms minimizes discrepancies between model predictions and observations, optimizing model fit to the data [Hastie et al., 2009]. Third, testing of machine learning algorithms enables direct evaluation of performance metrics, such as power and accuracy, on a dataset that is independent of that used for training [Hastie et al., 2009]. Fourth, machine learning algorithms are tailored to making predictions from data representing many correlated or conflicting features of varying levels of importance [Hastie et al., 2009], which is a critical consideration when using gene expression data from multiple conditions and species. Finally, CLOUD demonstrates high power and accuracy in predicting both expression divergence and its evolutionary parameters of duplicate genes in two species [DeGiorgio and Assis, 2021], suggesting that taking a similar approach with single-copy genes may yield favorable performance as well.

Thus, PiXi employs an adaptation of the multi-layer neural network of CLOUD [DeGiorgio and Assis, 2021], as well as two additional machine learning architectures—random forest and support vector machine—to account for different linear and nonlinear relationships in the input data. Specifically, PiXi uses each machine learning architecture to classify the expression of single-copy genes in two species as either “conserved” or “diverged”, and to estimate the parameters underlying their evolution. Application of PiXi to simulated data shows that all of its machine learning architectures have high power and accuracy in predicting both expression divergence and parameters across a wide range of evolutionary scenarios, with the multi-layer neural network globally outperforming other architectures. Moreover, application of PiXi to empirical data in *Drosophila* reveals that approximately 20% of positionally relocated genes undergo expression divergence driven by a combination of neutral and selective forces, and that such genes are often involved in cellular energy production. PiXi has been implemented as an open source R package, which is available at http://assisgroup.fau.edu/software.html and https://github.com/rassis/PiXi. Input data can include gene expression measurements in a single or multiple conditions, making PiXi applicable to studying expression divergence in both single- and multicellular organisms.

## Results

### Construction of PiXi

PiXi is constructed on an OU model of gene expression evolution [Hansen, 1997, Butler and King, 2004, Kalinka et al., 2010, Brawand et al., 2011, Perry et al., 2012, Rohlfs et al., 2014, Rohlfs and Nielsen, 2015, DeGiorgio and Assis, 2021]. In particular, suppose we have gene expression data from multiple conditions for single-copy orthologous genes in two species, Species 1 and Species 2. We model the expression evolution of these genes along the phylogeny relating the two species as an OU process, in which expression is pulled toward optimal states *θ*_1_ in Species 1 and *θ*_2_ in Species 2 through selection with strength *α*, and randomly fluctuates through phenotypic drift with strength *σ*^2^. In this study, we consider two scenarios for the optimal expression states in Species 1 and Species 2: *θ*_1_ = *θ*_2_, which should result in “conserved” gene expression between the species, and *θ*_1_ ≠ *θ*_2_, which should result in “diverged” gene expression between the species.

Following Brawand et al. [2011], gene expression in the two species **e** = (*e*_1_, *e*_2_) under this OU process is distributed as multivariate normal with mean

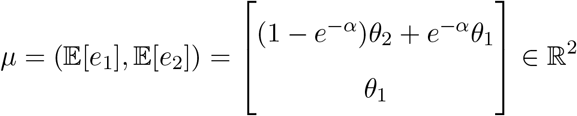

and covariance matrix

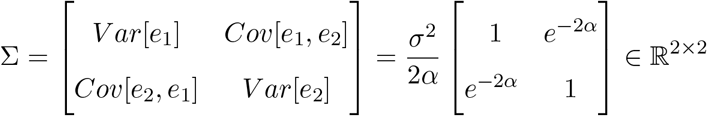

Though we assume here that gene expression is independent across conditions, this approach can be extended to account for an expression covariance structure [Revell and Harmon, 2008, Revell and Collar, 2009, Eastman et al., 2011, Clavel et al., 2015].

Here we consider the problem in which genes in the two species can be split into two sets: a sample set 𝒮 that contains the genes of interest, and a background set ℬ that contains other genes in the genome. We therefore let

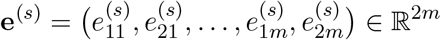

and

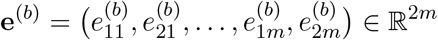

be the expression vectors for sample gene *s* ∈ 𝒮 and background gene *b* ∈ ℬ, where 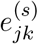 and 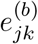 are respectively the expression levels for sample gene *s* and background gene *b* in species *j* ∈ {1, 2} and condition *k* ∈ {1, 2, …, *m*}.

Following DeGiorgio and Assis [2021], we transform and compare the expression vector **e**^(*s*)^ of each sample gene *s* ∈ 𝒮 to the expression vector **e**^(*b*)^ of each background gene *b* ∈ ℬ to obtain the input feature vector

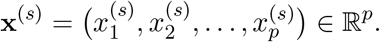

Specifically, the input feature vector **x**^(*s*)^ contains a set of *p* = 2*m* + 21 derived features (Table 1), many of which utilize comparisons to distributions of Euclidean distances dist(ℬ) and Pearson correlation coefficients cor(ℬ) between expression vectors of background genes ℬ in the two species. Comparison to this background set is amenable in a machine learning framework, but would not be in maximum likelihood methods based on OU models.

**Table 1:**
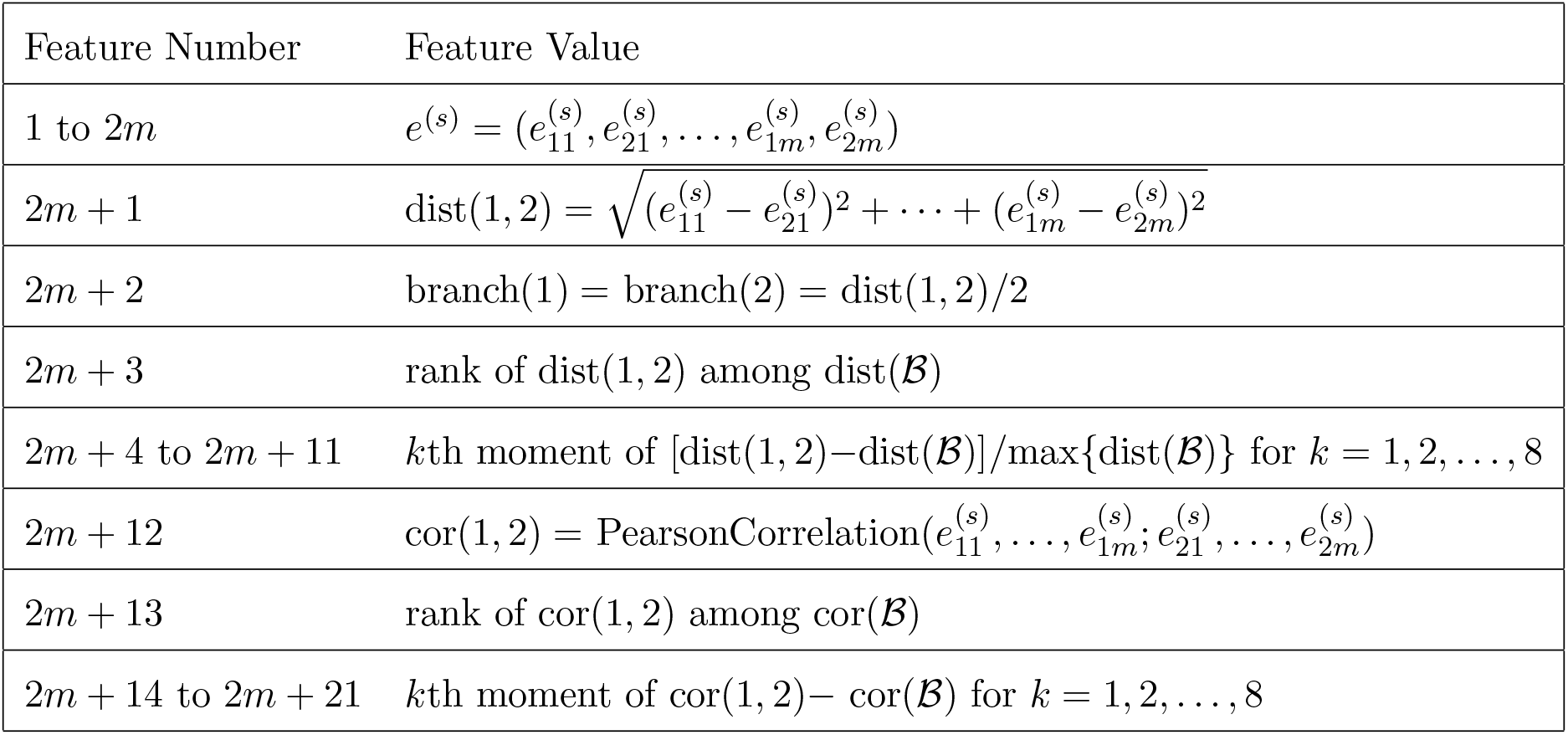
Set of *p* = 2*m* + 21 derived features used as input to `PiXi`

| Feature Number | Feature Value |
| --- | --- |
| 1 to $2m$ | $e^{(s)} = (e_{11}^{(s)}, e_{21}^{(s)}, \dots, e_{1m}^{(s)}, e_{2m}^{(s)})$ |
| $2m + 1$ | $\text{dist}(1, 2) = \sqrt{(e_{11}^{(s)} - e_{21}^{(s)})^2 + \dots + (e_{1m}^{(s)} - e_{2m}^{(s)})^2}$ |
| $2m + 2$ | $\text{branch}(1) = \text{branch}(2) = \text{dist}(1, 2)/2$ |
| $2m + 3$ | rank of $\text{dist}(1, 2)$ among $\text{dist}(\mathcal{B})$ |
| $2m + 4$ to $2m + 11$ | $k$th moment of $[\text{dist}(1, 2) - \text{dist}(\mathcal{B})]/\max\{\text{dist}(\mathcal{B})\}$ for $k = 1, 2, \dots, 8$ |
| $2m + 12$ | $\text{cor}(1, 2) = \text{PearsonCorrelation}(e_{11}^{(s)}, \dots, e_{1m}^{(s)}; e_{21}^{(s)}, \dots, e_{2m}^{(s)})$ |
| $2m + 13$ | rank of $\text{cor}(1, 2)$ among $\text{cor}(\mathcal{B})$ |
| $2m + 14$ to $2m + 21$ | $k$th moment of $\text{cor}(1, 2) - \text{cor}(\mathcal{B})$ for $k = 1, 2, \dots, 8$ |

Given the input feature vector **x**^(*s*)^, we seek to predict the output response *y*^(*s*)^. When performing classification to predict expression divergence of sample gene *s* ∈ 𝒮, *y*^(*s*)^ is the label for *K* = 2 classes “conserved” and “diverged”. In contrast, when performing regression to predict evolutionary parameters of sample gene *s* ∈ 𝒮, *y*^(*s*)^ is the quantitative response for *K* = 4*m* parameter estimates in each of the *m* conditions, where in each condition we obtain parameter estimates for optimal expression states *θ*_1_ and *θ*_2_, strength of selection *α*, and strength of phenotypic drift *σ*^2^. To account for a diversity of linear and nonlinear relationships, we implement three machine learning architectures for performing these classification and regression tasks: multi-layer neural network (NN), random forest (RF), and support vector machine (SVM, see *Methods*).

### Prediction performance of PiXi on simulated data

To evaluate the prediction performance of PiXi, we trained and tested its three machine learning architectures on independent balanced datasets of sample genes 𝒮 simulated under “conserved” and “diverged” expression classes (see *Methods*). The training set consisted of 20,000 observations (10,000 for each class), and the test set consisted of 2,000 observations (1,000 for each class). Evolutionary parameters for each dataset were drawn independently and uniformly at random across many orders of magnitude, with *θ*_1_, *θ*_2_ ∈ [0, 5], log_10_(*α*) ∈ [0, 3], and log_10_(*σ*^2^) ∈ [−2, 3] for each of *m* = 6 conditions, matching the number of tissues in an empirical dataset on which we later applied PiXi (see *Application of PiXi to empirical data in Drosophila*). This yielded a total of 24 random parameters per simulated replicate, as well as *p* = 2*m* + 21 = 33 derived features (Table 1) used for training the NN, RF, and SVM. We trained and tested the three machine learning architectures of PiXi on these datasets to enable direct comparisons of their performance.

We first assessed performance of the NN, RF, and SVM architectures of PiXi in classifying gene expression as either “conserved” or “diverged”. For comparison, we also followed previous studies in constructing another expression distance-based classifier [Assis and Bachtrog, 2013, Perry and Assis, 2016], using five-fold cross-validation to select a cutoff for defining expression divergence with this classifier (see *Methods*). Analysis of the resulting classifications reveals that all machine learning architectures of PiXi outperform the distance-based classifier, with the best overall performance achieved by a NN composing two hidden layers (Figure 1; see *Methods*). In particular, across the wide parameter space explored, classification power is highest with the NN, substantially lower with the RF and SVM, and lowest with the distance-based classifier (Figure 1A). Similarly, classification accuracy is approximately 92.75% with the NN, 79.75% with the RF, 80.45% with the SVM, and 77.95% with the distance-based classifier. Further, all machine learning architectures of PiXi exhibit more balanced classification rates than the distance-based classifier, with the highest balance observed with the NN (Figure 1B). Specifically, correct predictions of the two classes are approximately 92.4% and 93.1% with the NN, 75.0% and 84.5% with the RF, 77.7% and 83.2% with the SVM, and 89.5% and 66.4% with the distance-based classifier (main diagonals of Figure 1B). Thus, the NN is not skewed in its class predictions, the RF and SVM are both skewed toward the “diverged” class, and the distance-based classifier is skewed toward the “conserved” class.

**Figure 1:**
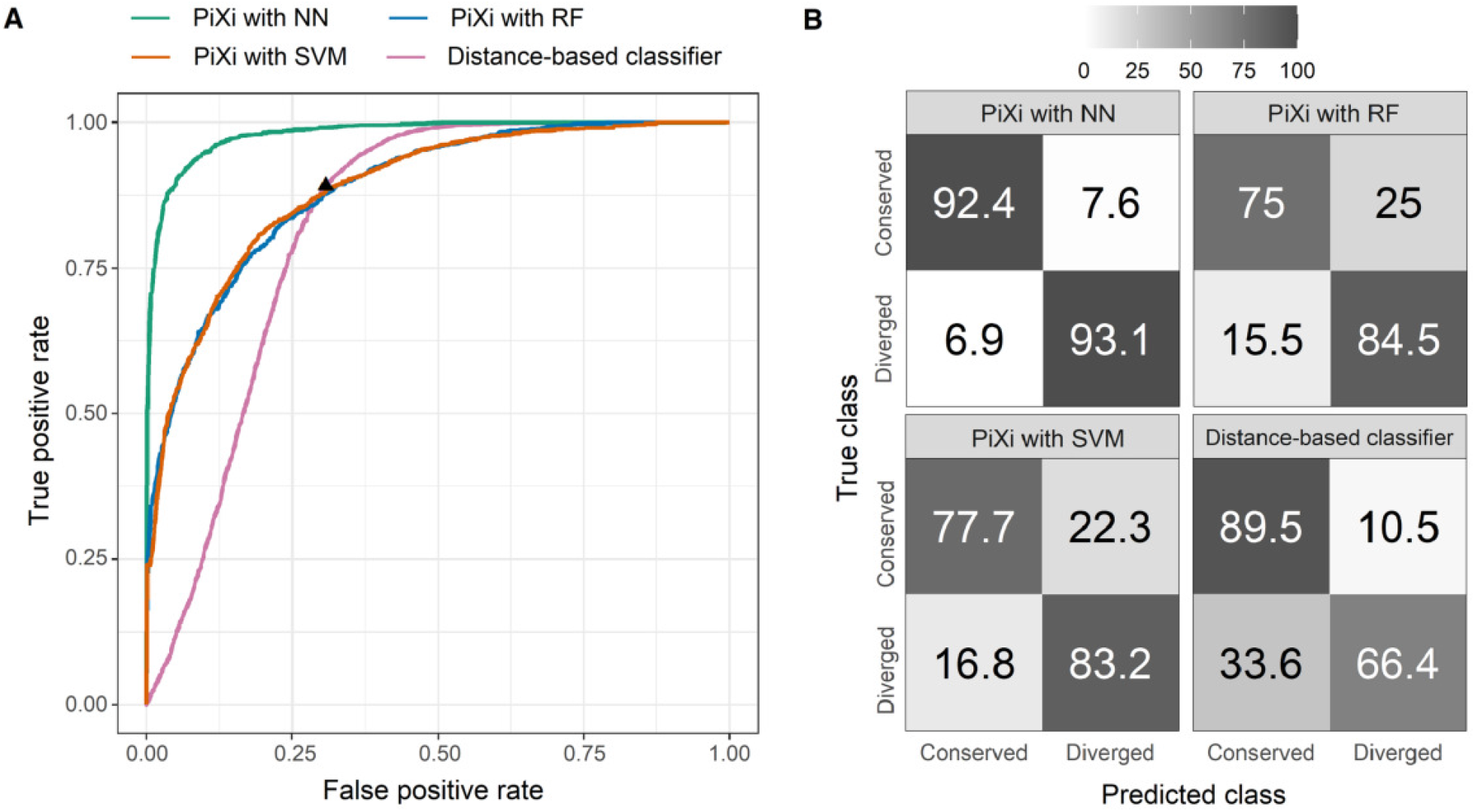
Classification performance of three machine learning architectures of `PiXi` and a distance-based classifier applied to test data simulated under parameters *α* ∈ [1, 10^3^] and *σ*^2^ ∈ [10^−2^, 10^3^] for each of the two classes. (*A*) Receiver operating characteristic curves showing the power of each method across the full range of false positive rates, with a black triangle depicting the cutoff chosen by cross-validation for the distance-based classifier. (*B*) Confusion matrices depicting classification rates of the two classes for each method.

Additionally, we find that the classification performance of all methods varies across smaller regions of the parameter space with combinations of strengths of selection (*α*) and phenotypic drift (*σ*^2^) representing specific evolutionary scenarios (Figures S1-S6). In general, the methods have higher classification power and accuracy when selection is strong (large *α*) or phenotypic drift is weak (small *σ*^2^), and lower classification power and accuracy when selection is weak (small *α*) or phenotypic drift is strong (large *σ*^2^). However, even under evolutionary scenarios for which classification is difficult (small *α* or small *σ*^2^), all machine learning architectures of PiXi still have substantially higher power and accuracy than the distance-based classifier, consistent with previous findings for the CLOUD predictor of duplicate gene expression divergence [DeGiorgio and Assis, 2021]. However, in contrast to our findings when considering the entire parameter space, all machine learning architectures show comparable classification performance when the parameter space is restricted, with similar classification power and accuracy for each combination of *α* and *σ*^2^ examined. This may be due to similarities in values of features across conditions when test data derive from a limited parameter space. Further, all machine learning architectures of PiXi produce balanced classification rates for every region of the parameter space, whereas the distance-based classifier appears to be swayed by phenotypic drift, preferentially choosing “conserved” when it is weak (small *σ*^2^) and “diverged” when it is strong (large *σ*^2^).

Aside from improved classification performance relative to a distance-based classifier, a major advantage of the machine learning framework of PiXi is its ability to predict parameters underlying gene expression evolution. Hence, we next assessed the parameter prediction accuracy of each of the machine learning architectures of PiXi on the same dataset used for classification. In particular, we were interested in how well PiXi could predict the four parameters in our OU model of gene expression evolution: optimal expression states *θ*_1_ and *θ*_2_, strength of selection *α*, and strength of phenotypic drift *σ*^2^. To compare parameter prediction accuracy among the machine learning architectures of PiXi, as well as between class labels, we examined distributions of mean prediction errors computed across the six tissues (Figure 2). This analysis reveals that all machine learning architectures yield unbiased parameter estimates, with mean prediction errors centered on zero. Additionally, consistent with CLOUD [DeGiorgio and Assis, 2021], predictions of *θ*_1_ and *θ*_2_ are more precise than those of *α* and *σ*^2^. Further, predictions of *θ*_1_ and *θ*_2_ are more precise for the “conserved” class, likely due to the additional degree of freedom in estimating parameters for the “diverged” class. Despite these general trends, the NN globally outperforms the RF and SVM architectures in parameter prediction, in that it displays the highest precision for both classes.

**Figure 2:**
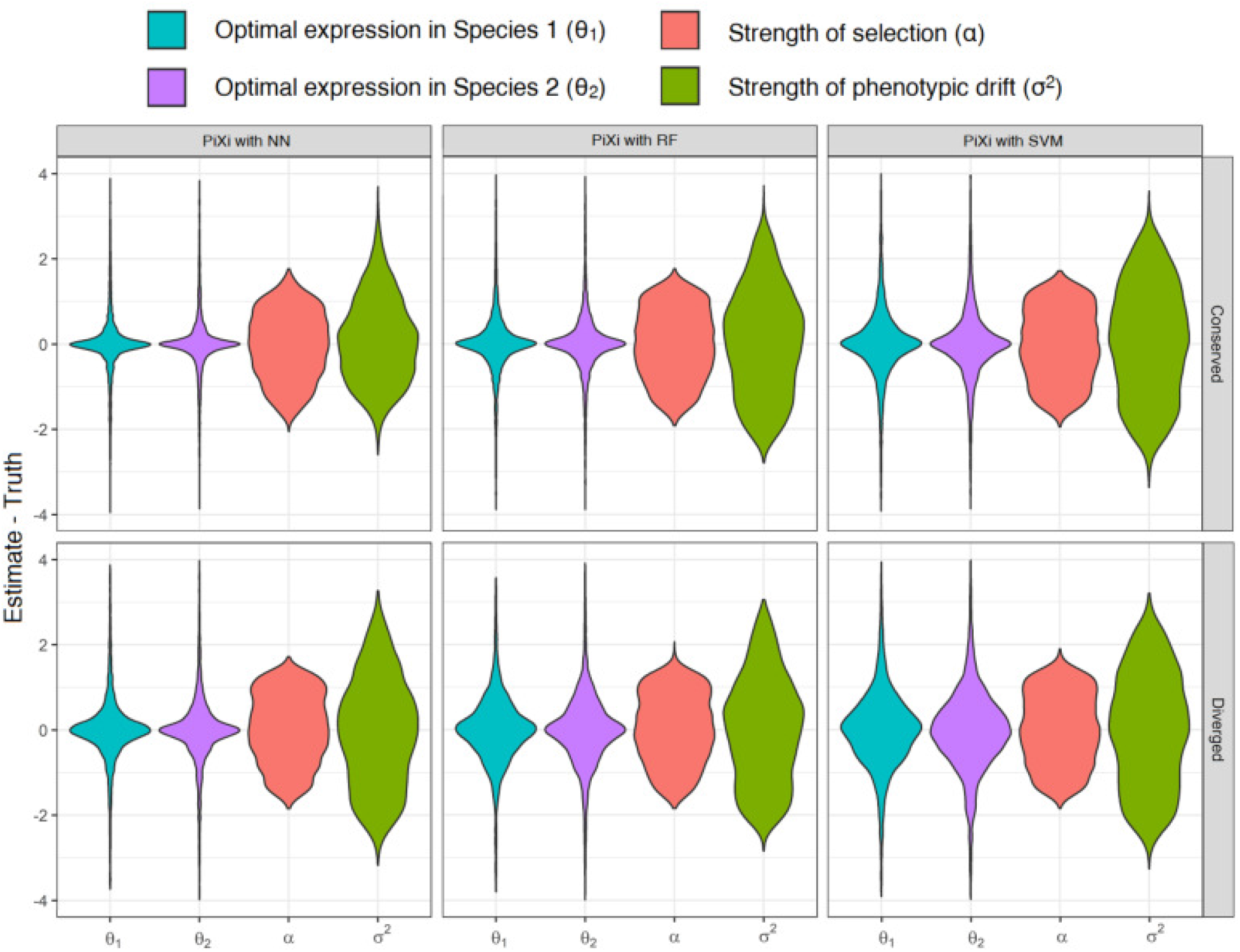
Parameter prediction performance of three machine learning architectures of `PiXi` applied to test data simulated under parameters *α* ∈ [1, 10^3^] and *σ*^2^ ∈ [10^−2^, 10^3^] for each of the two classes. Violin plots display distributions of mean parameter prediction errors across the *m* = 6 conditions for each simulated test dataset.

As with classification, parameter prediction accuracy of all machine learning architectures of PiXi varies similarly across smaller regions of the parameter space representing specific evolutionary scenarios (Figures S7-S9). Specifically, though these methods always produce unbiased estimates of *θ*_1_ and *θ*_2_, their estimates of *α* and *σ*^2^ are biased toward higher or lower values in some regions of the parameter space. Moreover, all parameter estimates tend to be more precise when selection is strong (large *α*) or phenotypic drift is weak (small *σ*^2^), and less precise when selection is weak (small *α*) or phenotypic drift is strong (large *σ*^2^). These findings mirror those observed with CLOUD [DeGiorgio and Assis, 2021]. Finally, whereas all machine learning architectures demonstrate comparable performance in predicting parameters in most evolutionary scenarios, the NN slightly outperforms the others in some instances, generally displaying less bias when phenotypic drift is strong (large *σ*^2^) and more precision for estimates of *θ*_1_ and *θ*_2_ when phenotypic drift is weak (small *σ*^2^).

### Application of PiXi to empirical data from *Drosophila*

Our simulation experiments demonstrate that PiXi has high power and accuracy in predicting both gene expression divergence and evolutionary parameters of single-copy genes in two species, and also that optimal performance is achieved through utilization of a NN architecture with two hidden layers (see *Methods*). Thus, we next applied PiXi with the same two-layer NN architecture to predict expression divergence and evolutionary parameters of 102 positionally relocated single-copy genes in two species of *Drosophila* [Hart et al., 2018] from their expression data in six tissues [Assis, 2019a] (see *Methods*). We chose this dataset because positional relocations may lead to expression divergence by introducing genes to new chromatin environments, which strongly influence their expression patterns and functions [Kleinjan and van Heyningen, 1998, Cohen et al., 2000, Boutanaev et al., 2002, Lercher et al., 2003, Hurst et al., 2004, Williams and Bowles, 2004, Michalak, 2008, Weber and Hurst, 2011, Assis, 2016]. The positional relocations in this dataset occurred between chromosomal arms and were polarized, with 53 and 49 inferred to have relocated in the *D. melanogaster* and *D. pseudoobscura* lineages, respectively [Hart et al., 2018]. Hence, to enable comparisons of optimal expression states before and after positional relocations, we set “Species 1” as the species with the gene on the ancestral chromosomal arm and optimal expression state *θ*_1_, and “Species 2” as the species with the gene on the derived chromosomal arm and optimal expression state *θ*_2_.

Of the 102 positionally relocated genes in our empirical dataset, 20 were classified as “diverged” by PiXi (Table S1). Moreover, examinations of distributions of parameter estimates reveal several clear distinctions between “conserved” and “diverged” classes (Figure 3). For one, estimates of *θ*_1_ and *θ*_2_ are larger for the “diverged” class, suggesting that optimal gene expression levels are higher for genes that underwent expression divergence after positional relocation in *Drosophila*. Second, estimates of *θ*_1_ and *θ*_2_ are similar for the “conserved” class and different for the “diverged” class, consistent with expectations under these two class scenarios of our OU model (see *Construction of PiXi*). Third, estimates of *θ*_2_ are generally larger than those of *θ*_1_ for the “diverged” class, indicating that optimal gene expression levels are higher for genes on derived chromosomal arms. Fourth, estimates of selection strength *α* and phenotypic drift *σ*^2^ are not significantly different between “conserved” and “diverged” genes, indicating that similar evolutionary forces act on all positionally relocated genes in *Drosophila*. Finally, while their distributions are not significantly different between classes, estimates of *α* and *σ*^2^ are both more precise for the “diverged” class. Thus, it appears that strengths of selection and phenotypic drift may both be more tightly controlled in genes that undergo expression divergence after positional relocation in *Drosophila*.

**Figure 3:**
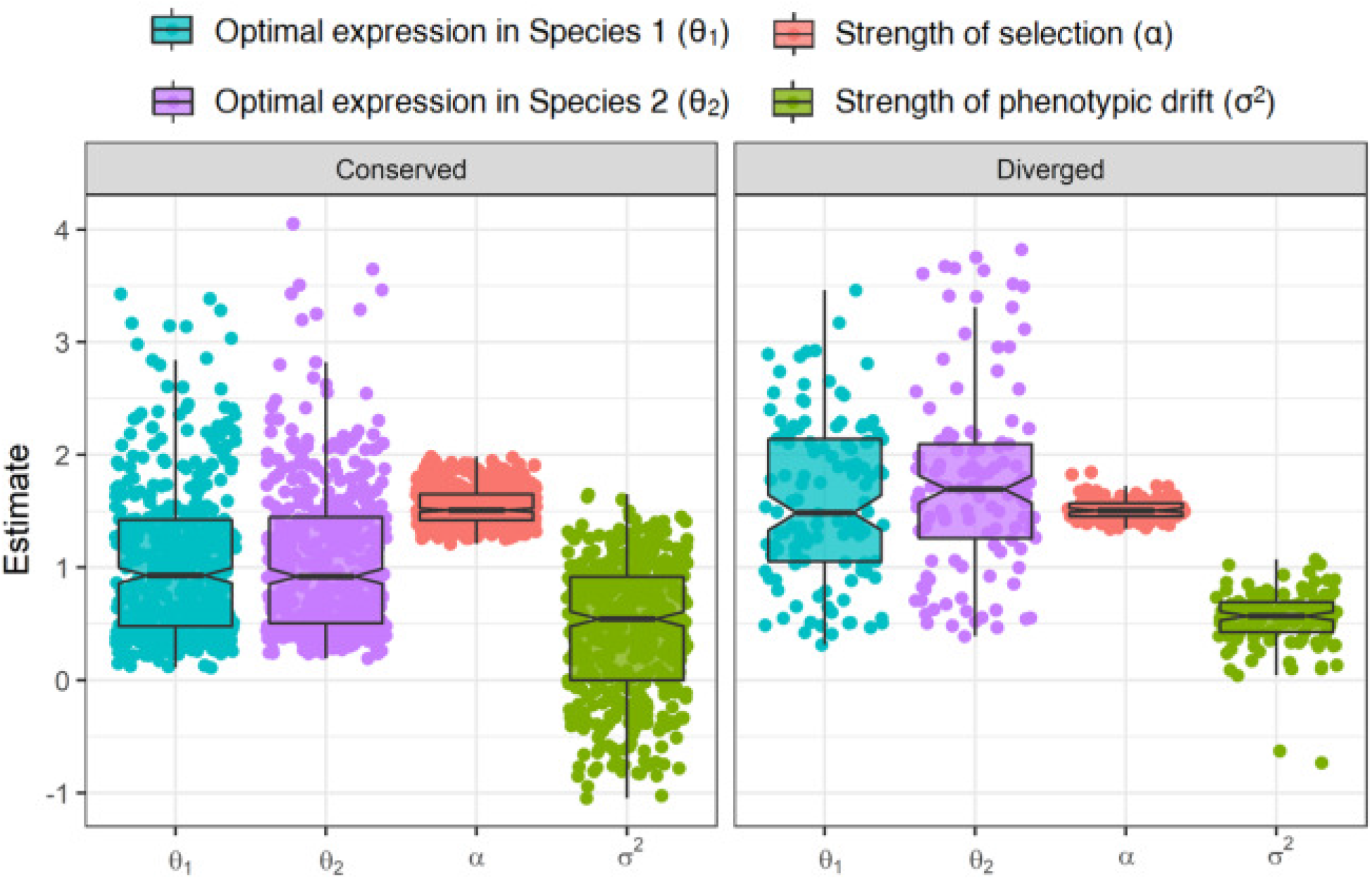
Parameter prediction results from application of the neural network architecture of `PiXi` to empirical data from positionally relocated genes in *Drosophila* [Hart et al., 2018, Assis, 2019a]. Box plots overlaid onto strip plots show distributions of log-transformed parameter estimates for each class. Note that six estimates, corresponding to the six tissues in the empirical dataset, are plotted for each parameter.

Investigations of the 20 *Drosophila* genes in the “diverged” class did not reveal any significant biases in the lineage in which positional relocations occurred (*P* = 0.51, binomial test), in either ancestral or derived chromosomal arm distributions (*P* = 0.79 and *P* = 0.86, respectively, Fisher’s exact tests), or in movements between X chromosomes and autosomes (*P* = 0.29, Fisher’s exact test), relative to expectations based on frequencies in the original dataset [Hart et al., 2018] (see *Methods*). However, it is important to note that the small sample size of the “diverged” class may limit our power to detect such biases. To better understand the biological factors that may contribute to gene expression divergence after positional relocation in *Drosophila*, we also analyzed functional annotations of genes classified as “conserved” and “diverged” (see *Methods*). Unfortunately, no results were statistically significant after multiple testing corrections, again perhaps as a result of the small sample size. Yet, several genes in the “conserved” class are involved in regulation of transcription and post-translational modifications in the nucleus (Table S2). In contrast, a few genes in the “diverged” class participate in the electron transport chain in the mitochondrial membrane, and particularly in the processes of ubiquinol cytochrome c reductase activity and oxidative phosphorylation (Table S3). This distinction illustrates that the functions of genes may dictate their evolutionary fates after positional relocations. Specifically, perhaps cellular energy production is more malleable than transcription and translation in *Drosophila*, and genes with such functions are therefore more likely to experience divergence after positional relocations.

For further analysis, we performed a case study of the *UQCR-11L* gene (Table S1; FBgn0050354 in *D. melanogaster*, FBgn0086842 in *D. pseudoobscura*) in the “diverged” class. We chose this gene, as it demonstrated the largest difference between optimal expression states *θ*_1_ and *θ*_2_, strongest selection *α*, and weakest phenotypic drift *σ*^2^ in the “diverged” class. *UQCR-11L*, or Ubiquinol-cytochrome c reductase 11 kDa subunit-like, underwent a positional relocation from the Muller E chromosomal arm to the Muller C chromosomal arm in the *D. melanogaster* lineage. Intriguingly, a previous study revealed that the positional relocation of *UQCR-11L* in the *D. melanogaster* lineage resulted in its insertion into the intron of another gene, *Acsl*, or Acyl-CoA synthetase long-chain [Assis, 2016]. Due to transcriptional interference, such “nested” genes were found to experience rapid sequence and expression divergence [Assis, 2016], consistent with our classification of *UQCR-11L* expression as “diverged”. Our parameter predictions also show that rapid sequence and expression divergence after this nesting event were likely driven by a combination of relatively strong selection (*α* ≈ 41.69) and moderate phenotypic drift (*σ*^2^ ≈ 1.55). Further, *UQCR-11L* is one of the handful of genes from our functional annotation analysis that participate in ubiquinol cytochrome c reductase activity in mitochondrial electron transport. Thus, *UQCR-11L* represents an interesting example for which positional relocation resulted in gene nesting, rapid sequence and expression divergence likely driven by strong selection against transcriptional interference, and perhaps corresponding functional divergence altering cellular energy production in *D. melanogaster*.

## Discussion

In this work, we present PiXi, an OU model-based machine learning framework for predicting expression divergence and its evolutionary parameters between single-copy genes in two species. PiXi implements three machine learning architectures for its predictions: NN, RF, and SVM. We demonstrate that each of these machine learning architectures has high power and accuracy in discriminating between “conserved” and “diverged” expression classes (Figures 1 and S1-S5), as well as high accuracy in estimating evolutionary parameters (Figures 2 and S7-S9), with the overall best performance for both tasks achieved by the NN. Moreover, these three machine learning architectures all globally outperform an expression distance-based classifier, which has the lowest classification power, accuracy, and balance (Figures 1 and S1-S6), as well as an inability to predict the evolutionary parameters underlying expression evolution. Hence, PiXi represents a significant advancement for the widespread problem of assaying expression divergence and its evolutionary parameters in a set of single-copy genes from two species. Though here we focused on usage with gene expression data from multiple conditions, PiXi can also be employed with gene expression data from a single condition, enabling its application to studies of gene expression divergence in both single- and multicellular organisms.

We chose to incorporate NN, RF, and SVM machine learning architectures in PiXi to allow for different types of linear and nonlinear relationships, as well as for variation in other properties, of the input data. In particular, the NN is linear when the number of hidden layers *L* = 0 and nonlinear otherwise, the RF is always nonlinear, and the SVM behaves as linear when the *γ* hyperparameter of its RBF kernel is small and as nonlinear otherwise. Further, though the NN outperformed the other architectures in our study, the RF and SVM architectures may be advantageous for properties of input data that we did not consider. For example, the RF may be beneficial if expression data are absent for some genes or conditions due to its robustness to missing data, whereas the SVM may be beneficial if expression data are measured in one or few conditions due to its ability to expand the dimensionality of the data. Thus, we kept all three machine learning architectures in the final version of PiXi to provide users with the flexibility to choose an architecture that is best suited to their data.

We also considered instead performing class and parameter predictions with a maximum likelihood framework, which has been used for other studies of expression evolution with OU models [Kalinka et al., 2010, Brawand et al., 2011, Perry et al., 2012, Rohlfs et al., 2014, Rohlfs and Nielsen, 2015]. Specifically, given expression data for Species 1 and Species 2, one can use maximum likelihood to estimate the set of parameters {*θ*_1_, *θ*_2_, *α, σ*^2^} from an OU model of expression evolution for the two classes, with constraints *θ*_1_ = *θ*_2_ for the “conserved” class and *θ*_1_ ≠ *θ*_2_ for the “diverged” class. Then one can employ a likelihood ratio test to discriminate between classes, with the “conserved” class representing the null hypothesis and the “diverged” class representing the alternative hypothesis. However, there are a few obstacles to this approach. For one, it would be highly dependent on underlying model assumptions, such as independence among conditions. Second, the “diverged” class, which has four free parameters per condition, would be over-parameterized without the inclusion of a third gene from an outgroup species. Third, it would be difficult to integrate the comparison to the background set of genes. Hence, we believe that using machine learning for predictions is ideal for the particular evolutionary problem at hand. As an empirical study, we applied the best-performing NN architecture of PiXi to expression data [Assis, 2019a] from 102 positionally relocated single-copy genes in two species of *Drosophila* [Hart et al., 2018]. Of these genes, 20 were classified as “diverged” (Table S1), supporting the hypothesis that movement of genes to new chromatin environments can lead to modification of their expression profiles. There were also some interesting distinctions between parameter estimates of “conserved” and “diverged” genes (Figure 3), together suggesting that genes that undergo expression divergence tend to have higher optimal expression levels before relocation, even higher optimal expression levels after relocation, and tighter control of selection and phenotypic drift than genes that retain their ancestral expression. Our follow-up analyses also revealed that several “conserved genes” are involved in transcriptional and post-transcriptional regulation (Table S2), whereas several “diverged” genes are involved in the electron transport chain (Table S3), perhaps indicating that expression divergence tends to impact cellular energy production. Further, our case study of the “diverged” gene with the largest difference between optimal expression states *θ*_1_ and *θ*_2_, strongest selection *α*, and weakest drift *σ*^2^ revealed it to be one of the handful of genes that participate in the electron transport chain, as well as a “nested” gene that relocated into an intron of another gene. Hence, our empirical study illustrates that application of PiXi can yield novel and interesting insights into the evolutionary trajectories and forces acting on single-copy genes.

## Methods

### Design of NN, RF, and SVM architectures for PiXi

In constructing the NN architecture for PiXi, we follow the approach of DeGiorgio and Assis [2021], tailoring it to our problem where appropriate. In particular, we consider a dense feed-forward neural network with *L* ∈ {0, 1, 2, 3} hidden layers, in which the first hidden layer has *p*[1] = 256 hidden units, and hidden layer *ℓ* ∈ {1, 2, …, *L*} has *p*[*ℓ*] = 256*/*2^*ℓ*−1^ hidden units, such that each hidden layer contains half the number of hidden units as the previous hidden layer [DeGiorgio and Assis, 2021]. To simplify our notation, we set the input layer as hidden layer zero, such that *p*[0] = *p* = 2*m* + 21 is the number of input features, and the output layer as hidden layer *L* + 1, such that *p*[*L* + 1] = *K*. The values at unit *k* ∈ {1, 2, …, *p*[*ℓ*]} of hidden layer *ℓ* ∈ {0, 1, 2, …, *L*} for sample gene *s* ∈ 𝒮 are defined by its activation 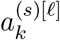. Because hidden layer zero is the input layer and hidden layer *L* + 1 is the output layer, the activation is given by

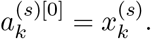

We seek to predict the output response vector

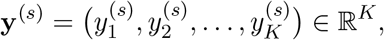

where

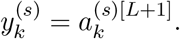

Continuing to follow the approach of DeGiorgio and Assis [2021], we define the activation for unit *k* of hidden layer *ℓ* ∈ {1, 2, …, *L*} as a nonlinear transformation of the linear combination of the activations for the previous hidden layers. Specifically, we apply the rectified linear unit [ReLU, Goodfellow et al., 2016] function defined as ReLU(*x*) = max(0, *x*), such that the activation for unit *k* in hidden layer *ℓ* of sample gene *s* is

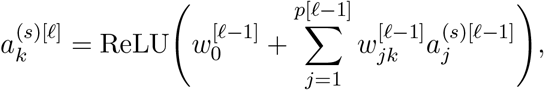

where 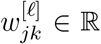 is the weight (parameter) from unit *j* in layer *ℓ* to unit *k* in layer *ℓ* + 1, and 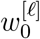 is the bias for layer *ℓ* [Goodfellow et al., 2016]. The output layer takes inputs from layer *L*, and has a different form depending on whether we consider the classification or the regression problem. For classification, we use the softmax activation function [Goodfellow et al., 2016], such that the output for class *k* ∈ {1, 2} of sample gene *s* is the probability

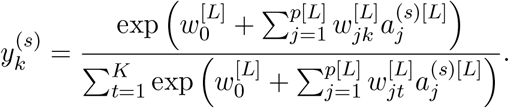

For regression, we use the linear activation function [Goodfellow et al., 2016], such that the output for parameter prediction *k* ∈ {1, 2, …, 4*m*} of sample gene *s* is

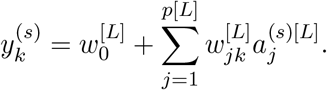

When *L* = 0, the NN simplifies to a linear model with multinomial regression for the classification problem and to linear regression for the regression problem [Hastie et al., 2009].

In designing the RF architecture for PiXi, we implement Breiman’s algorithm [Breiman, 2001] with *p* = 2*m* + 21 features and *n* = 500 trees. RF is an ensemble learner that makes predictions from a “forest” of *n* randomly constructed trees [Breiman, 2001]. To construct each tree in the random forest, a bootstrap training set of 20,000 observations is created through random sampling with replacement from the 20,000 observations in the original training set. Then, for each split in the tree, a subset of size 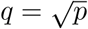 of the features is selected uniformly at random [Wright and Ziegler, 2017], and the node is split on of these *q* features by minimizing node impurity, which is computed with the Gini index [Gini, 1936] for classification and the estimated response variances [Wright et al., 2017] for regression. The tree is grown without pruning [Breiman, 2001], with a minimum node size of ten for classification and five for regression. This process is repeated to construct each of the 500 trees in the forest [Breiman, 2001]. For classification, each tree contains estimated class probabilities [Malley et al., 2012], and the output class *k* ∈ {1, 2} of sample gene *s* ∈ *S* is chosen as the class with the larger mean estimated probability across the 500 trees [Breiman, 2001]. For regression, the output parameter prediction *k* ∈ {1, 2, …, 2*m*} of sample gene *s* ∈ *S* is given by the mean parameter estimate across the 500 trees [Breiman, 2001].

In developing the SVM architecture for PiXi, we use a radial basis function (RBF) kernel [Hastie et al., 2009] of form

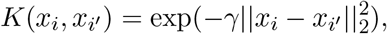

with *p* = 2*m* + 21 features and 11 *γ* ∈ [0.001, 5] hyperparameters uniformly chosen on a logarithmic scale. Though the RBF kernel is nonlinear, it behaves as a linear kernel when *γ* is small [Hastie et al., 2009], thereby enabling us to capture both linear and nonlinear relationships in the input data. Using this kernel to transform the feature space, the SVM identifies the maximum margin hyperplane [Hastie et al., 2009] defined by *x* ∈ ℝ^*p*^ such that

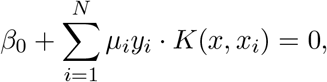

where *β*_0_ is the intercept and *µ*_1_, *µ*_2_, …, *µ*_*N*_ are the coefficients of the support vectors (*i*.*e*., those *x*_*i*_ with *µ*_*i*_ *>* 0) in the Lagrange dual function that maximize the margin, or the distance between training observations and the hyperplane [Hastie et al., 2009].

For classification, the maximum margin hyperplane results in optimal separation of classes [Cortes and Vapnik, 1995], and the output class *k* ∈ {1, 2} of sample gene *s* ∈ *S* is selected based on the sign of *y*^(*s*)^, which specifies on which side of the hyperplane it lies. For regression, the maximum margin hyperplane results in optimal fit to the training data [Drucker et al., 1997], with the margin in this case representing the maximum unpenalized residual *ϵ*, or difference between observed and predicted parameter *k* ∈ {1, 2, …, 4*m*}. The predicted parameter *k* ∈ {1, 2, …, 4*m*} of sample gene *s* ∈ *S* is given by the value of 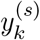.

All described machine learning architectures were implemented in R (2021). We used Keras [Chollet et al., 2017] with a TensorFlow backend [Abadi et al., 2015] for the NN, ranger [Wright and Ziegler, 2017] for the RF, and liquidSVM [Steinwart and Thomann, 2017] for the SVM. Note that when training the regression models, the NN was allowed to jointly estimate all *K* = 4*m* model parameters, whereas a separate regression was performed for each parameter within the RF and SVM frameworks.

### Training PiXi on simulated data

To train the three machine learning architectures, we first generated a balanced simulated dataset with *N* = 20, 000 training observations, 10, 000 from each of the two classes. We assumed independence among tissues, and that there were a total of *m* = 6 tissues as in an empirical gene expression dataset from *Drosophila* [Assis, 2019a] on which we later applied our method (see *Application of PiXi to empirical data in Drosophila*), for a total of *p* = 33 input features. To ensure that the simulated dataset was realistic, we drew model parameters *θ*_1_, *θ*_2_ ∈ [0, 5] to match the range observed in the empirical *Drosophila* expression data [Assis, 2019a], and *α* from log_10_(*α*) ∈ [0, 3] and *σ*^2^ from log_10_(*σ*^2^) ∈ [−2, 3] to consider a wide range of potential strengths for selection and phenotypic drift. The class *k* was determined to be “conserved” when *θ*_1_ = *θ*_2_ and “diverged” when *θ*_1_ ≠ *θ*_2_. Then, we simulated gene expression data **e**^(*i*)^ ∈ ℝ^2*m*^ under model parameters for a given class *k*, generating *N*_*k*_ simulated replicates of parameter values.

To train the NN, we followed DeGiorgio and Assis [2021] by minimizing the elastic net [Zou and Hastie, 2005] penalized cost function

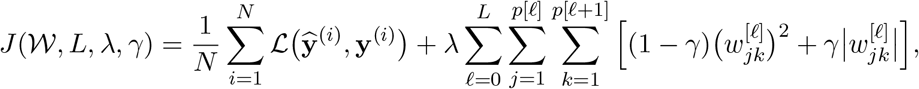

where 𝒲 is the set of parameter estimates, *L* is the number of hidden layers, *λ* is a tuning parameter that reduces the complexity of the fitted model by shrinking the weights to zero, *γ* ∈ [0, 1] is a tuning parameter that determines the influence of the *L*_1_- and *L*_2_-norm penalties for simultaneous feature selection, and 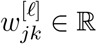 is the weight (parameter) from unit *j* in layer *ℓ* to unit *k* in layer *ℓ*+1. As in DeGiorgio and Assis [2021], we estimated the set of parameters 𝒲 from a number of hidden layers *L* on the pair of regularization tuning parameters *λ* and *γ* using the Adam optimizer [Kingma and Ba, 2014] with learning rate 10^−3^ and exponential decay rates for the first and second moment estimates of *β*_1_ = 0.9 and *β*_2_ = 0.999 [Kingma and Ba, 2014]. Similarly, we also used mini-batch optimization with a batch size of 5,000 observations for 500 epochs, and five-fold cross-validation [Hastie et al., 2009] to estimate *L, λ*, and *γ* [DeGiorgio and Assis, 2021]. In particular, here we used 16,000 (80%) observations for training, with the remaining 4,000 (20%) held out for validation. We also balanced each sample dataset, with equal numbers of observations from each class in the training (8,000) and validation (2,000) sets. Following DeGiorgio and Assis [2021], we considered values of *L* ∈ {0, 1, 2, 3} and *γ* ∈ {0, 0, 0.1, …, 1.0}, as well as 25 values of *λ* chosen uniformly across log_10_(*λ*) ∈ [−12, −3]. Given the optimal cross-validation estimates 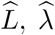, and 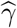 for *L, λ*, and *γ*, respectively, we estimated the neural network model parameters 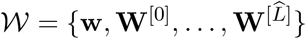 using all 20,000 training observations. Consistent with the findings of DeGiorgio and Assis [2021], a neural network with 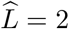 hidden layers provided the best cross-validation performance, with a validation loss for classification of approximately 0.191 with optimal tuning parameters 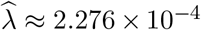 and 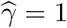, and a validation loss for regression of approximately 0.504 with optimal tuning parameters 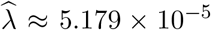 and 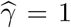. These values of 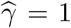 imply that the *L*_1_-norm penalty was solely employed by our elastic net regularization in both the classification and regression settings, which encouraged sparse models with maximal feature selection.

To train the RF, we performed bagging [Breiman, 1996] in tandem with random feature selection, as described by Breiman [2001]. In particular, a bootstrap sample training set consisting of 20,000 observations was constructed through random sampling with replacement from the 20,000 observations in the original sample training set. Due to bootstrapping, approximately 1*/*3 of observations in the original training set were left out [Efron, 1979]. We used the bootstrap sample to build a random forest with *n* = 500 trees to predict classes and evolutionary parameters. Each tree in the random forest was grown such that on every split, we let the tree choose among the 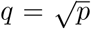 features that minimize node impurity, with a minimum node size of ten for classification and five for regression.

To train the SVM, we maximized the Lagrangian dual function [Hastie et al., 2009]

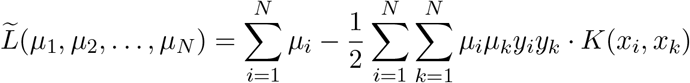

subject to the constraint

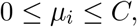

where *µ*_1_, *µ*_2_, …, *µ*_*N*_ are the dual function parameters that maximize the margin *M* of the support vectors (*x*_*i*_ with *µ*_*i*_ *>* 0), *K*(*x*_*i*_, *x*_*k*_) is the RBF kernel function with hyperparameter *γ* that influences the width of the kernel function, and *C* is a tuning parameter that defines penalization of observations within *M*. As with our NN, we used five-fold cross-validation [Hastie et al., 2009] to estimate *γ* and *C*, again with 16,000 (80%) observations for training and the remaining 4,000 (20%) held out for validation. Similarly, we balanced each dataset, with equal numbers of observations from each class in the training (8,000) and validation (2,000) sets.

### Testing PiXi on simulated data

The training dataset for sample genes 𝒮 was simulated from an OU process, which is described in subsection *Training machine learning architectures on data simulated from OU processes* above. However, in that subsection, we assumed that the training dataset for background genes ℬ was given. To obtain this dataset, we followed DeGiorgio and Assis [2021], using a Brownian motion model [Felsenstein, 1973] to generate 10,000 six-tissue expression vectors inspired by those of single-copy non-relocated genes in *Drosophila* [Meisel et al., 2009], such that our trained models are based on the typical level of expression divergence observed. All machine learning architectures of PiXi were trained on these training sample 𝒮 and background ℬ training datasets.

After model training, we evaluated the performance of the three machine learning architectures of PiXi on an independent balanced test dataset of 2,000 simulated observations, 1,000 from each of the two classes. As when generating our training dataset, we assumed *m* = 6 independent tissues and drew OU model parameters uniformly at random, with *θ*_1_, *θ*_2_ ∈ [0, 5] to match the range observed in the empirical *Drosophila* expression data [Assis, 2019a], and *α* from log_10_(*α*) ∈ [0, 3] and *σ*^2^ from log_10_(*σ*^2^) ∈ [−2, 3] to consider a wide range of potential strengths for selection and phenotypic drift. The class *k* was determined to be “conserved” when *θ*_1_ = *θ*_2_ and “diverged” when *θ*_1_ ≠ *θ*_2_, and gene expression data were generated **e**^(*i*)^ ∈ ℝ^2*m*^ under model parameters for a given class *k*, resulting in 1,000 simulated replicates of parameter values.

We also examined the accuracy of each machine learning architecture of PiXi on test datasets drawn from restricted regions of the parameter space. In particular, we used the same approach outlined above to simulate test datasets of 2,000 obseverations, 1,000 from each class, for three distinct ranges of *α* ∈ [1, 10], [10, 100], and [100, 1, 000], and five distinct ranges of *σ*^2^ ∈ [0.01, 0.1], [0.1, 1], [1, 10], [10, 100], and [100, 1, 000]. For each combination of a range of *α* and a range of *σ*^2^, we sampled *α* and *σ*^2^ uniformly at random.

For evaluation of the classification performance of these machine learning architectures, we constructed another distance-based classifier with a cutoff *c* for selecting the output class *k*. In particular, we first computed Euclidean and Manhattan distances between absolute and relative expression levels across *m* = 6 conditions in the training dataset for background genes ℬ that was used by the machine learning architectures. For each of these four sets of distances, we uniformly selected 100 cutoff values from the range of distances, and used five-fold cross-validation to select the value of *c* that maximized accuracy. Then, we constructed four classifiers, each with a different distance metric and optimal value of *c*. We compared the power and accuracy of these four classifiers by applying them to the test dataset that we used for the three machine learning architectures. Of these distance-based classifiers, the classifier with Manhattan distances between absolute expression levels and with *c* ≈ 7.26 selected by cross-validation had the highest power and accuracy (Figure S10). Thus, we used this best distance-based classifier for comparisons with the three machine learning architectures of PiXi.

### Analysis of empirical data from *Drosophila*

We applied PiXi with the two-layer neural network architecture that demonstrated optimal performance (see *Testing machine learning architectures on data simulated from OU processes*) to empirical data consisting of positionally relocated and non-relocated single-copy genes in *D. melanogaster* and *D. pseudoobscura* [Hart et al., 2018] and their expression abundances measured in the same six tissues from each species [Assis, 2019a]. To produce this input dataset, we first obtained 127 positionally relocated and 7,977 non-relocated single-copy genes in *D. melanogaster* and *D. pseudoobscura* from Hart et al. [2018]. Hart et al. [2018] identified positionally relocated single-copy genes through curation of previously annotated inter-chromosomal-arm positional relocations that occurred along the lineages leading to *D. melanogaster* and *D. pseudoobscura* [Hahn et al., 2007, Meisel et al., 2009], and inferred their ancestral and derived chromosomal arms through comparisons to the chromsomal arms of their orthologs in *D. willistoni, D. virilis*, and *D. grimshawi* genomes. In that study, Hart et al. [2018] also defined non-relocated single-copy genes as those retained as 1:1:1:1:1 orthologs on the same chromosomal arm in all five *Drosophila* species [Meisel et al., 2009].

Next, we obtained quantile-normalized gene expression abundances for carcass, female head, ovary, male head, testis, and accessory gland tissues in *D. melanogaster* and D. pseudoobscura from the Dryad dataset associated with Assis [2019a] at https://doi.org/10.5061/dryad.742564m. Briefly, Assis [2019a] downloaded paired-end RNA-sequencing reads from modENCODE [Celniker et al., 2009] at https://www.modencode.com, aligned these reads to the reference transcriptomes of each species with Bowtie 2 [Langmead et al., 2009], computed expression abundances of genes in fragments per kilobase of exon per million fragments mapped (FPKM) [Trapnell et al., 2013] with eXpress [Roberts and Pachter, 2013], and quantile-normalized and log-transformed these FPKM values in R (2021). We removed all Hart et al. [2018] genes for which the Assis [2019a] quantile-normalized FPKM *<* 1 in all six tissues for either *D. melanogaster* or *D. pseudoobscura*, yielding 102 positionally relocated and 6,820 non-relocated single-copy genes and corresponding gene expression abundances on which we applied PiXi.

We trained PiXi with a two-layer neural network architecture through five-fold cross-validation [Hastie et al., 2009], setting the regularization tuning parameters as 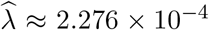 and 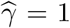 for classification, and 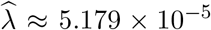 and 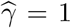 for regression (see *Training machine learning architectures on data simulated from OU processes*). Then, we applied the trained model to the 102 positionally relocated (sample set 𝒮) and 6,820 non-relocated (background set ℬ) single-copy genes in *Drosophila* to predict the expression of positionally relocated genes as either “conserved” or “diverged”, as well as their evolutionary parameters *θ*_1_, *θ*_2_, *α*, and *σ*^2^.

We used the DAVID Functional Annotation Tool [Huang et al., 2009a,b] to assay functions of genes classified as “conserved” and “diverged”. Specifically, we ran this tool twice, each time using the list of *D. melanogaster* genes from either the “conserved” or “diverged” predicted class as our gene list, and all other genes in the *D. melanogaster* genome as the background list. We also assessed lineage-specific biases in the “diverged” class with a two-tailed exact binomial test, in which we set the number of successes *x* = 12 to represent the number of “diverged” genes that underwent positional relocations in the *D. melanogaster* lineage, the number of trials *n* = 20 to represent the total number of “diverged” genes, and the probability of success *p* = 53*/*102 to represent the expected frequency of “diverged” genes that underwent positional relocations in the *D. melanogaster* lineage if it is equal to the total frequency of positional relocations in this lineage. Finally, we assayed biases in ancestral and derived chromosomal arm distributions, as well as in relocations between sex chromosomes and autosomes with two-tailed Fisher’s exact tests, in which we compared observed distributions of the “diverged” class to those expected based on their frequencies in the full dataset of positional relocations. All statistical analyses were performed in the R software environment [R Core Team, 2021].

## Data availability

The empirical data used in this article were obtained from sources in the public domain: modENCODE at https://www.modencode.com, and Dryad at https://doi.org/10.5061/dryad.742564m.

## Acknowledgments

This work was supported by National Institutes of Health grant R35GM142438 to RA, National Institutes of Health grant R35GM128590 to MD, National Science Foundation grant DEB-2001059 to RA, National Science Foundation grants DEB-1949268 and BCS-2001063 to MD, and National Science Foundation grant DBI-2130666 to RA and MD.

## Notes

### Competing Interest Statement

The authors have declared no competing interest.

## References

M Abadi, A Agarwal, P Barham, E Brevdo, Z Chen, C Citro, GS Corrado, A Davis, J Dean, M Devin, S Ghemawat, I Goodfellow, A Harp, G Irving, M Isard, Y Jia, R Jozefowicz, L Kaiser, M Kudlur, J Levenberg, D Mané, R Monga, S Moore, D Murray, C Olah, M Schuster, J Shlens, B Steiner, I Sutskever, K Talwar, P Tucker, V Vanhoucke, V Vasudevan, F Viégas, O Vinyals, P Warden, M Wattenberg, M Wicke, Y Yu, and Z Zheng. TensorFlow: Large-scale machine learning on heterogeneous systems. 2015. URL https://www.tensorflow.org/.

R Assis. Drosophila duplicate genes evolve new functions on the fly. Fly, 8:91–94, 2014.

R Assis. Transcriptional interference promotes rapid functional evolution of young Drosophil nested genes. Genome Biol Evol, 8:3149–3158, 2016.

R Assis. Out of the testis, into the ovary: biased outcomes of gene duplication and deletion in Drosophila. Evolution, 73:1850–1862, 2019a.

R Assis. Lineage-specific expression divergence in grasses is associated with male reproduction, host-pathogen defense, and domestication. Genome Biol Evol, 11:207–219, 2019b.

R Assis. No expression divergence despite transcriptional interference between nested protein-coding genes in mammals. Genes, 12:1381, 2021.

R Assis and D Bachtrog. Neofunctionalization of young duplicate genes in Drosophila. Proc Natl Acad Sci USA, 110:17409–17414, 2013.

R Assis and D Bachtrog. Rapid divergence and diversification of mammalian duplicate gene functions. BMC Evol Biol, 15:1–7, 2015.

R Assis and AS Kondrashov. Conserved proteins are fragile. Mol Biol Evol, 31:419–424, 2014.

R Assis, Q Zhou, and D Bachtrog. Sex-biased transcriptome evolution in Drosophila. Genome Biol Evol, 29:1189–1200, 2012.

N Bhardwaj and H Lu. Correlation between gene expression profiles and protein-protein interactions within and across genomes. Bioinf, 21:2730–2738, 2005.

G Blanc and KH Wolfe. Functional divergence of duplicated genes formed by polyploidy during Arabidopsis evolution. Plant Cell, 16:1679–1691, 2004.

AM Boutanaev, AI Kalmykova, YY Shevelyov, and DI Nurminsky. Large clusters of co-expressed genes in the Drosophila genome. Nature, 420:475–478, 2002.

D Brawand, M Soumillon, A Necsulea, P Julien, G Csárdi, P Harrigan, M Weier, A Liechti, A Aximu-Petri, M Kircher, and et al. The evolution of gene expression levels in mammalian organs. Nature, 478:343–348, 2011.

L Breiman. Bagging predictors. Machine Learning, 24:123–140, 1996.

L Breiman. Random forests. Machine Learning, 45:5–32, 2001.

MA Butler and AA King. Phylogenetic comparative analysis: a modeling approach for adaptive evolution. Am Nat, 164:683–695, 2004.

SB Carroll. Evolution at two levels: on genes and form. PLoS Biol, 3:e245, 2005.

SE Celniker, LA Dillon, MB Gerstein, KC Gunsalus, S Henikoff, GH Karpen, M Kellis, EC Lai, JD Lieb, DM MacAlpine, G Micklem, F Piano, M Snyder, L Stein, KP White, RH Waterston, and modENCODE Cosnortium. Unlocking the secrets of the genome. Nature, 459:927–930, 2009.

FJJ Chain, D Ilieva, and BJ Evans. Duplicate gene evolution and expression in the wake of vertebrate allopolyploidization. BMC Evol Biol, 8:1–16, 2008.

Allaire JJ Chollet, Fx et al. R interface to keras. 2017.

J Clavel, G Escarguel, and G Merceron. mvmorph: an r package for fitting multivariate evolutionary models to morphometric data. Methods Ecol Evol, 6:1311–1319, 2015.

BA Cohen, RD Mitra, JD Hughes, and GM Church. A computational analysis of whole-genome expression data reveals chromosomal domains of gene expression. Nat Genet, 26:183–186, 2000.

The ENCODE Project Consortium. An integrated encyclopedia of dna elements in the human genome. Nature, 489:57–74, 2012.

C Cortes and V Vapnik. Support-vector networks. Mach Learn, 20:273–297, 1995.

R De Smet, E Sabaghian, Z Li, Y Saeys, and Y Van de Peer. Coordinated functional divergence of genes after genome duplication in Arabidopsis thaliana. Plant Cell, 29:2786–2800, 2017.

M DeGiorgio and R Assis. Learning retention mechanisms and evolutionary parameters of duplicate genes from their expression data. Mol Biol Evol, 38:1209–1224, 2021.

H Drucker, CC Burges, L Kaufman, AJ Smola, and VN Vapnik. Support vector regression machines. Adv Neural Inf Process Syst, 9:155–161, 1997.

JM Eastman, ME Alfaro, P Joyce, AL Hipp, and LJ Harmon. A novel comparative method for identifying shifts in the rate of character evolution on trees. Evolution, 65:3578–3589, 2011.

B Efron. Bootstrap methods: another look at the jackknife. Ann Stat, 7:1–26, 1979.

J Felsenstein. Maximum-likelihood estimation of evolutionary trees from continuous characters. Am J Hum Genet, 25:471, 1973.

ZL Fuller, GD Haynes, S Richards, and SW Schaeffer. Genomics of natural populations: How differentially expressed genes shape the evolution of chromosomal inversions in Drosophila pseudoobscura. Genetics, 204:287–301, 2016.

C Gini. On the measure of concentration with special reference to income and statistics. Colorado College Publication, 208:73–79, 1936.

I Goodfellow, Y Bengio, and A Courville. Deep feedforward networks. Deep Learn, 2016.

X Gu. Statistical methods for testing functional divergence after gene duplication. Mol Biol Evol, 16:1664–1674, 1999.

X Gu. Maximum-likelihood approach for gene family evolution under functional divergence. Mol Biol Evol, 18:453–464, 2001.

MW Hahn, MV Han, and S-G Han. Gene family evolution across 12 Drosophila genomes. PLoS Genet, 3:e197, 2007.

TF Hansen. Stabilizing selection and the comparative analysis of adaptation. Evolution, 51: 1341–1351, 1997.

MLI Hart, BL Vu, Q Bolden, KT Chen, CL Oakes, L Zoronjic, and RP Meisel. Genes relocated between Drosophila chromosome arms evolve under relaxed selective constraints relative to non-relocated genes. J Mol Evol, 86:340–352, 2018.

T Hastie, R Tibshirani, and J Friedman. The elements of statistical learning: data mining, inference, and prediction. Springer, New York, NY, 2nd edition, 2009.

DW Huang, BT Sherman, and RA Lempicki. Systematic and integrative analysis of large gene lists using david bioinformatics resources. Nucleic Protoc, 4:44–57, 2009a.

DW Huang, BT Sherman, and RA Lempicki. Bioinformatics enrichment tools: paths toward the comprehensive functional analysis of large gene lists. Nucleic Acids Res, 37:1–13, 2009b.

BG Hunt, L Ometto, L Keller, and MAD Goodisman. Evolution at two levels in fire ants: the relationship between patterns of gene expression and protein sequence evolution. Mol Biol Evol, 30:263–271, 2012.

LD Hurst, C Pál, and MJ Lercher. The evolutionary dynamics of eukaryotic gene order. Nat Rev Genet, 5:299–310, 2004.

X Jiang and R Assis. Rapid functional divergence after small-scale duplication in grasses. BMC Evol Biol, 19:97, 2019.

X Jiang and R Assis. Genome-wide prediction, functional divergence, and characterization of stress-responsive bzr transcription factors in B. napus. Front Plant Sci, 4: doi.org/10.3389/fpls.2021.790655, 2022.

AT Kalinka, KM Varga, DT Gerrard, S Preibisch, DL Corcoran, J Jarrells, U Ohler, CM Bergman, and P Tomancak. Gene expression divergence recapitulates the developmental hourglass model. Nature, 468:811–814, 2010.

M Kapushesky, I Emam, E Holloway, P Kurnosov, Zorin A, and et al. Gene expression atlas at the european bioinformatics institute. Nucleic Acids Res, 38:D690–D698, 2010.

D Kingma and J Ba. Adam: a method for stochastic optimization. arXiv, 1412:6980, 2014.

DJ Kleinjan and V van Heyningen. Position effect in human genetic disease. Hum Mol Genets, 7: 1611–1618, 1998.

FA Kondrashov, IB Rogozin, YI Wolf, and EV Koonin. Selection in the evolution of gene duplications. Genome Biol, 3:1–9, 2002.

B Langmead, C Trapnell, M Pop, and SL Salzberg. Ultrafast and memory-efficient alignment of short dna sequences to the human genome. Genome Biol, 10:R25, 2009.

B Lemos, BR Bettencourt, CD Meiklejohn, and DL Hartl. Evolution of proteins and gene expression levels are coupled in Drosophila and are independently associated with mrna abundance, protein length, and number of protein-protein interactions. Mol Biol Evol, 22:1345–1354, 2005.

MJ Lercher, T Blumenthal, and LD Hurst. Coexpression of neighboring genes in Caenorhabditis elegans is mostly due to operons and duplicate genes. Genome Res, 13:238–243, 2003.

W-H Li, J Yang, and X Gu. Expression divergence between duplicate genes. Trends Genet, 21: 602–607, 2005.

N Lopez-Bigas, S De, and SA Teichmann. Functional protein divergence in the evolution of Homo sapiens. Genome Biol, 9:R33, 2008.

M Lynch and A Force. The probability of duplicate gene preservation by subfunctionalization. Genetics, 154:459–473, 2000.

VJ Lynch and GP Wagner. Resurrecting the role of transcription factor change in developmental evolution. Evolution, 62:2131–2154, 2008.

N Mähler, J Wang, BK Terebieniec, PK Ingvarsson, NR Street, and TR Hvidsten. Gene co-expression network connectivity is an important determinant of selective constraint. PLoS Genet, 13:e1006402, 2017.

KD Makova and W-H Li. Divergence in the spatial pattern of gene expression between human duplicate genes. Genome Res, 13:1638–1645, 2003.

JD Malley, J Kruppa, A Dasgupta, KG Malley, and A Ziegler. Probability machines: consistent probability estimation using nonparametric learning machines. Methods Inf Med, 51:74–81, 2012.

RP Meisel, MV Han, and MW Hahn. A complex suite of forces drives gene traffic from Drosophila x chromosomes. Genome Biol Evol, 1:176–188, 2009.

D Meng, Y Cao, T Chen, M Abdullah, Q Jin, and et al. Evolution and functional divergence of mads-box genes in Pyrus. Scientific Rep, 9:1266, 2019.

P Michalak. Coexpression, coregulation, and cofunctionality of neighboring genes in eukaryotic genomes. Genomics, 91:243–248, 2008.

BM Musungu, D Bhatnagar, RL Brown, GA Payne, G OBrian, AM Fakhoury, and M Geisler. A network approach of gene co-expression in the Zea mays/Aspergillus flavus pathosystem to map host/pathogen interaction pathways. Front Genet, 7:206, 2016.

NL Nehrt, WT Clark, P Radivojac, and MW Hahn. Testing the ortholog conjecture with comparative functional genomic data from mammals. PLoS Comp Biol, 7:e1002073, 2011.

SV Nuzhdin, ML Wayne, KL Harmon, and LM McIntyre. Common pattern of evolution of gene expression level and protein sequence in Drosophila. Mol Biol Evol, 21:1308–1317, 2004.

BR Perry and R Assis. Cdrom: Classification of duplicate gene retention mechanisms. BMC Evol Biol, 16:1–4, 2016.

GH Perry, P Melsted, JC Marioni, Y Wang, R Bainer, JK Pickrell, K Michelini, S Zehr, AD Yoder, M Stephens, and et al. Comparative rna sequencing reveals substantial genetic variation in endangered primates. Genome Res, 22:602–610, 2012.

R Petryszak, T Burdett, B Fiorelli, NA Fonseca, Gonzalez-Porta M, and et al. Expression atlas update—a database of gene and transcript expression from microarray- and sequencing-based functional genomics experiments. Nucleic Acids Res, 42:D926–D932, 2013.

R Core Team. R: A Language and Environment for Statistical Computing. R Foundation for Statistical Computing, Vienna, Austria, 2021. URL https://www.R-project.org/.

LJ Revell and DC Collar. Phylogenetic analysis of the evolutionary correlation using likelihood. Evolution, 63:1090–1100, 2009.

LJ Revell and LJ Harmon. Testing quantitative genetic hypotheses about the evolutionary rate matrix for continuous characters. Evol Ecol Res, 10:311–331, 2008.

A Roberts and L Pachter. Streaming fragment assignment for real-time analysis of sequencing experiments. Nat Methods, 10:71–73, 2013.

RV Rohlfs and R Nielsen. Phylogenetic anova: the expression variance and evolution model for quantitative trait evolution. Syst Biol, 64:695–708, 2015.

RV Rohlfs, P Harrigan, and R Nielsen. Modeling gene expression evolution with an extended ornstein-uhlenbeck process accounting for within-species variation. Mol Biol Evol, 31:201–211, 2014.

I Steinwart and P Thomann. liquidsvm: A fast and versatile svm package. arXiv preprint 1702.06899, 2017.

C Trapnell, DG Hendrickson, M Sauvageau, L Goff, JL Rinn, and L Pachter. Differential analysis of gene regulation at transcript resolution with rna-seq. Nat Biotechnol, 31:46–53, 2013.

CC Weber and LD Hurst. Support for multiple classes of local expression clusters in Drosophila melanogaster, but no evidence for gene order conservation. Genome Biol, 12:R23, 2011.

NE Wheeler, L Barquist, RA Kingsley, and PP Gardner. A profile-based method for identifying functional divergence of orthologous genes in bacterial genomes. Bioinf, 32:3566–3574, 2016.

EJB Williams and DJ Bowles. Coexpression of neighboring genes in the genome of Arabidopsis thaliana. Genome Res, 14:1060–1067, 2004.

GA Wray, MW Hahn, E Abouheif, JP Balhoff, M Pizer, MV Rockman, and LA Romano. The evolution of transcriptional regulation in eukaryotes. Mol Biol Evol, 20:1377–1419, 2003.

Marvin N. Wright and Andreas Ziegler. ranger: A fast implementation of random forests for high dimensional data in C++ and R. Journal of Statistical Software, 77(1):1–17, 2017. doi: 10.18637/jss.v077.i01.

MN Wright, T Dankowski, and A Ziegler. Unbiased split variable selection for random survival forests using maximally selected rank statistics. Stat Med, 36:1272–1284, 2017.

X Zhong, M Lundberg, and L Raberg. Divergence in coding sequence and expression of different functional categories of immune genes between two wild rodent species. Genome Biol Evol, 13: evab023, 2021.

H Zou and T Hastie. Regularization and variable selection via the elastic net. Stat Methodol, 67: 301–320, 2005.

