## Supplementary Figures for "Predicting expression divergence and its evolutionary parameters between single-copy genes in two species"

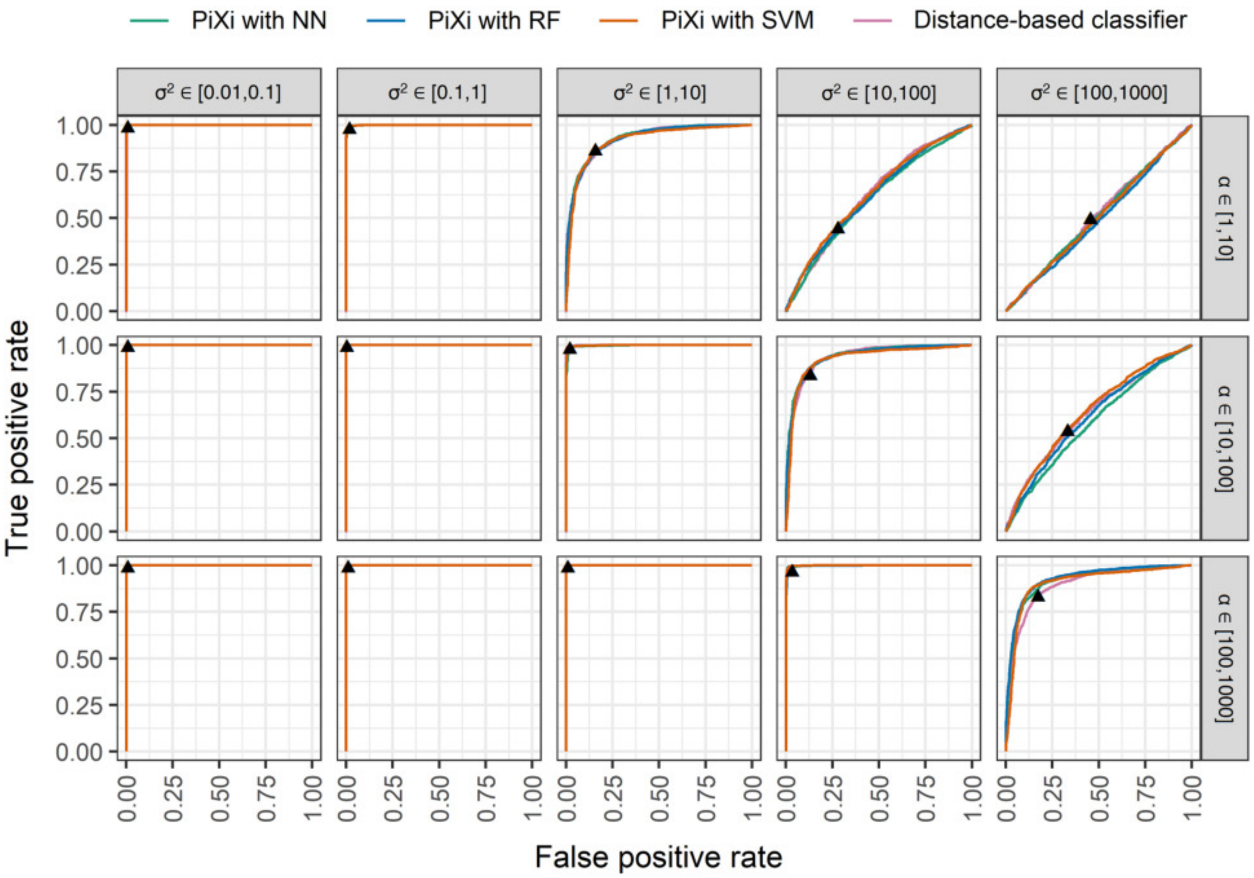


Figure S1: Receiver operating characteristic curves for the three machine learning architectures of `PiXi` and a distance-based classifier applied to test data simulated under specific parameter ranges for $\alpha$ and $\sigma^2$. Classification power is highest for large $\alpha$ and small $\sigma^2$, and lowest for small $\alpha$ and large $\sigma^2$. The black triangles depict the cutoff chosen by cross-validation for the distance-based classifiers.

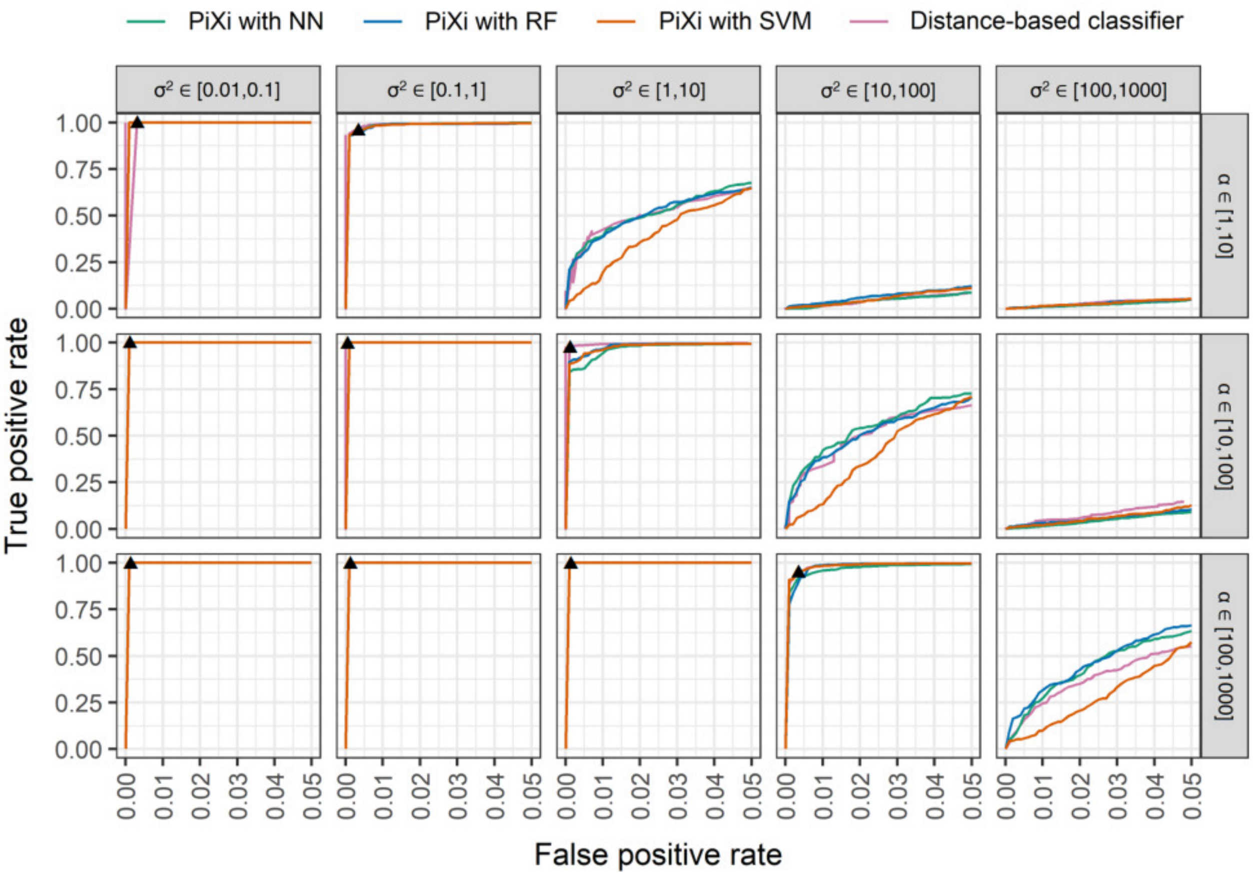


Figure S2: Receiver operating characteristic curves truncated at a false positive rate of 5% for the three three machine learning architectures of `PiXi` and a distance-based classifier applied to test data simulated under specific parameter ranges for $\alpha$ and $\sigma^2$. Classification power is highest for large $\alpha$ and small $\sigma^2$, and lowest for small $\alpha$ and large $\sigma^2$. The black triangles depict the cutoff chosen by cross-validation for the distance-based classifiers.

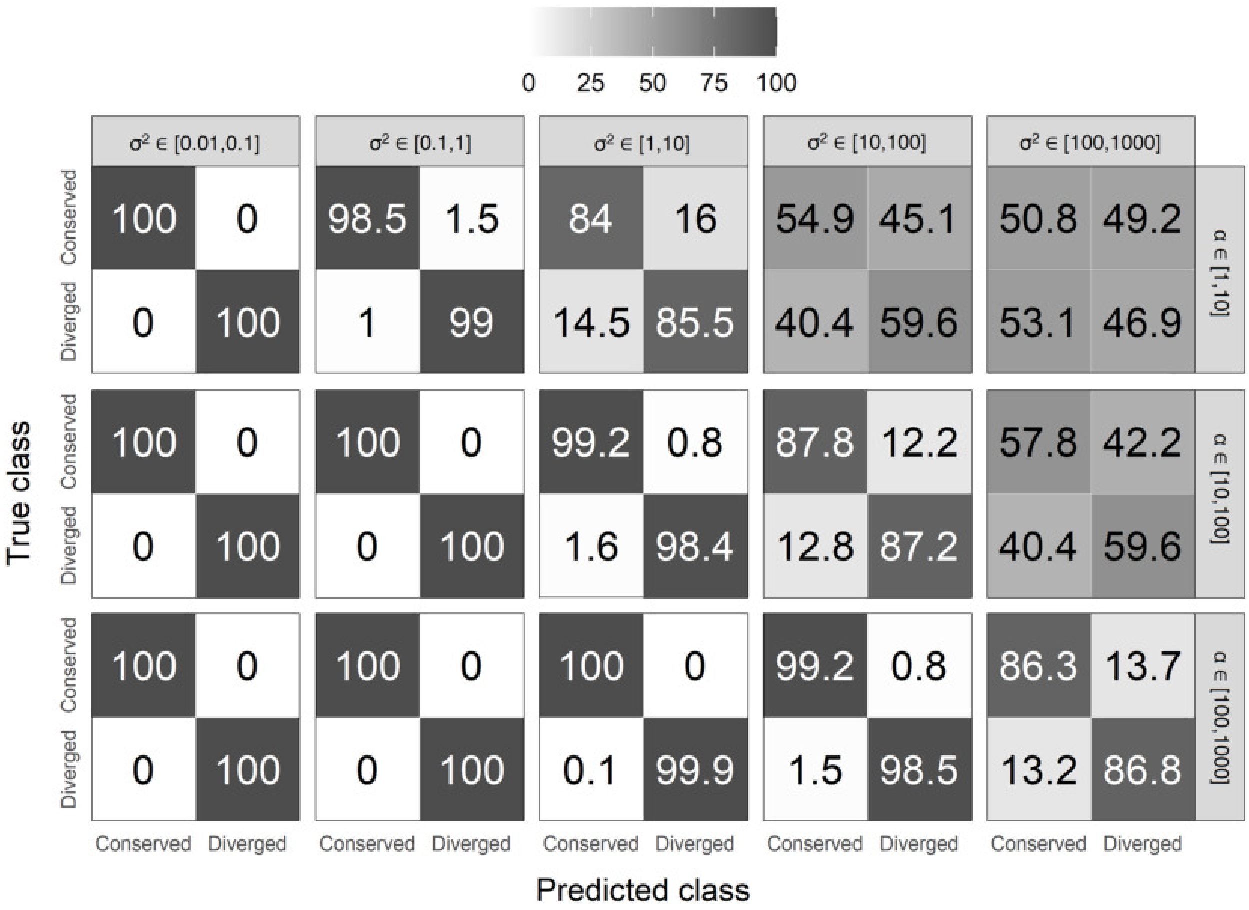


Figure S3: Confusion matrices depicting classification rates of the two classes for the neural network architecture of `PiXi` applied to test data simulated under specific parameter ranges for $\alpha$ and $\sigma^2$. Classification accuracy is highest for large $\alpha$ and small $\sigma^2$, and lowest for small $\alpha$ and large $\sigma^2$.

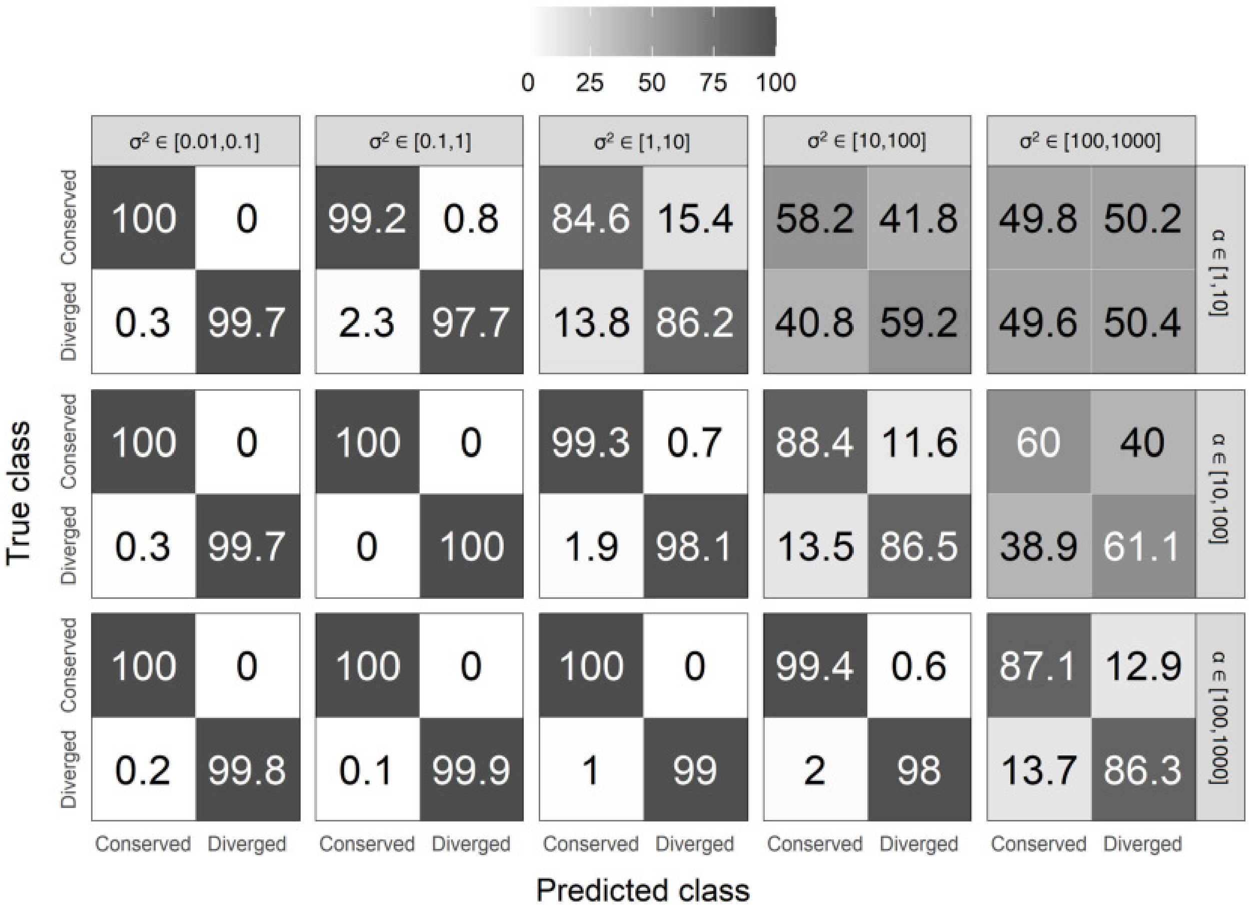


Figure S4: Confusion matrices depicting classification rates of the two classes for the random forest architecture of `PiXi` applied to test data simulated under specific parameter ranges for $\alpha$ and $\sigma^2$. Classification accuracy is highest for large $\alpha$ and small $\sigma^2$, and lowest for small $\alpha$ and large $\sigma^2$.

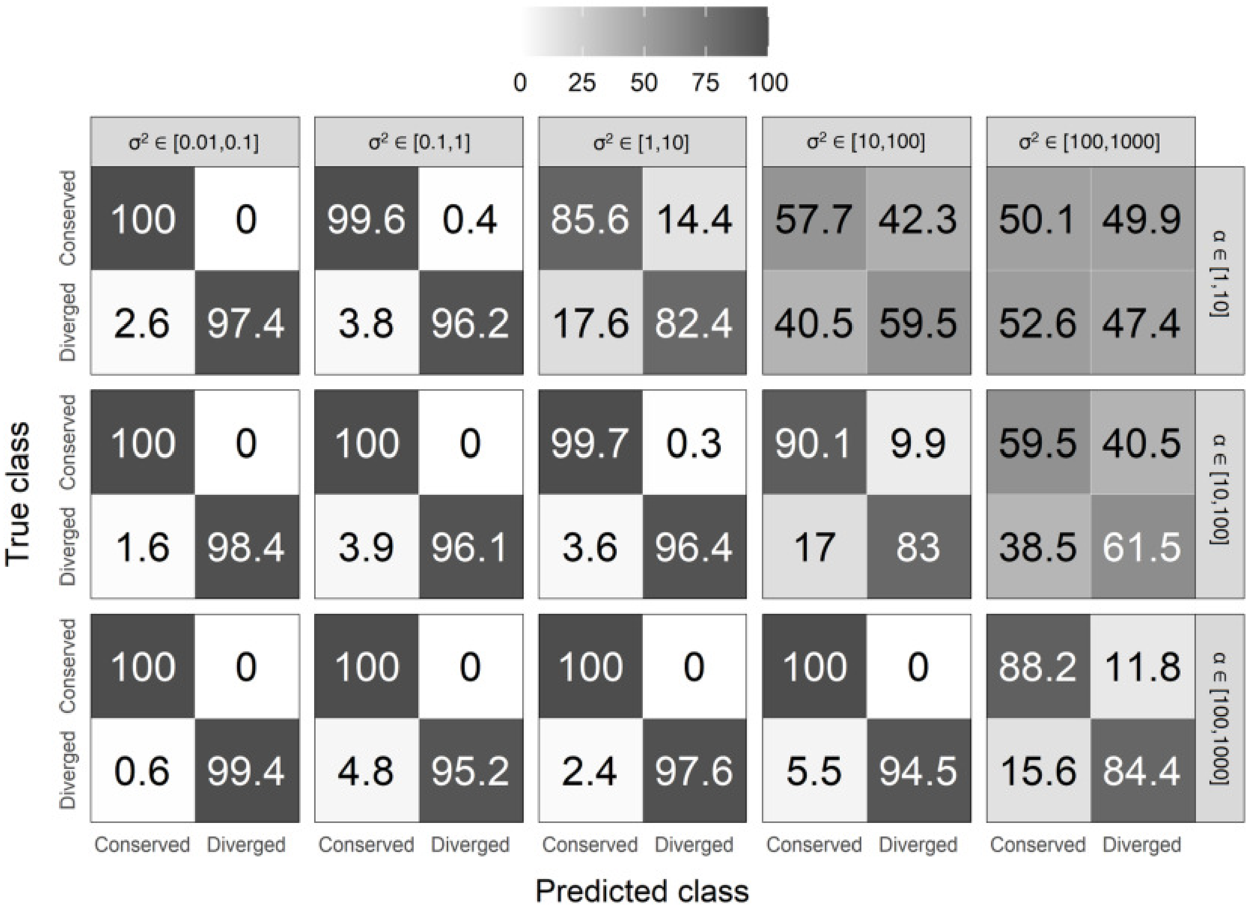


Figure S5: Confusion matrices depicting classification rates of the two classes for the support vector machine architecture of `PiXi` applied to test data simulated under specific parameter ranges for $\alpha$ and $\sigma^2$. Classification accuracy is highest for large $\alpha$ and small $\sigma^2$, and lowest for small $\alpha$ and large $\sigma^2$.

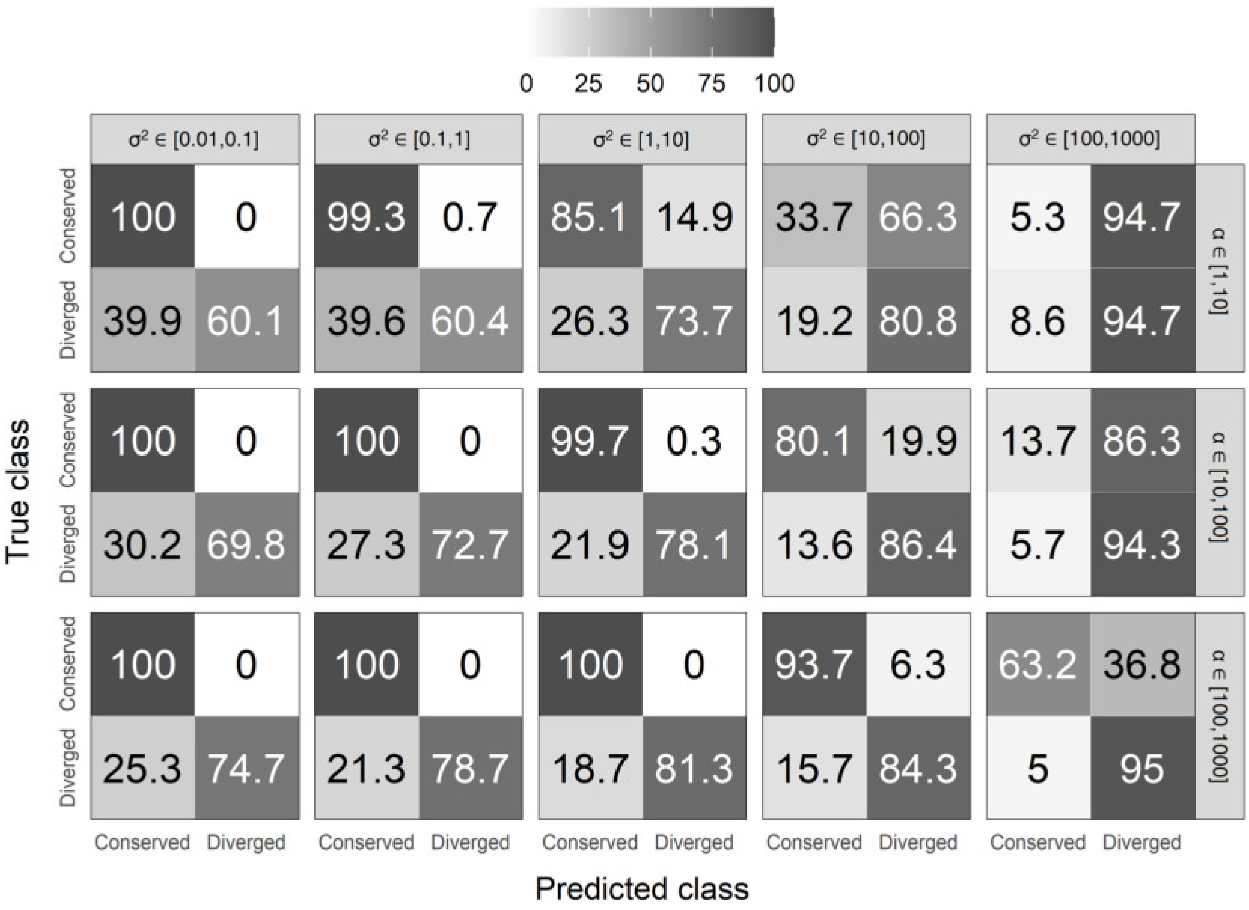


Figure S6: Confusion matrices depicting classification rates of the two classes for the distance-based classifier applied to test data simulated under specific parameter ranges for $\alpha$ and $\sigma^2$. Classification accuracy is highest for large $\alpha$ and small $\sigma^2$, and lowest for small $\alpha$ and large $\sigma^2$.

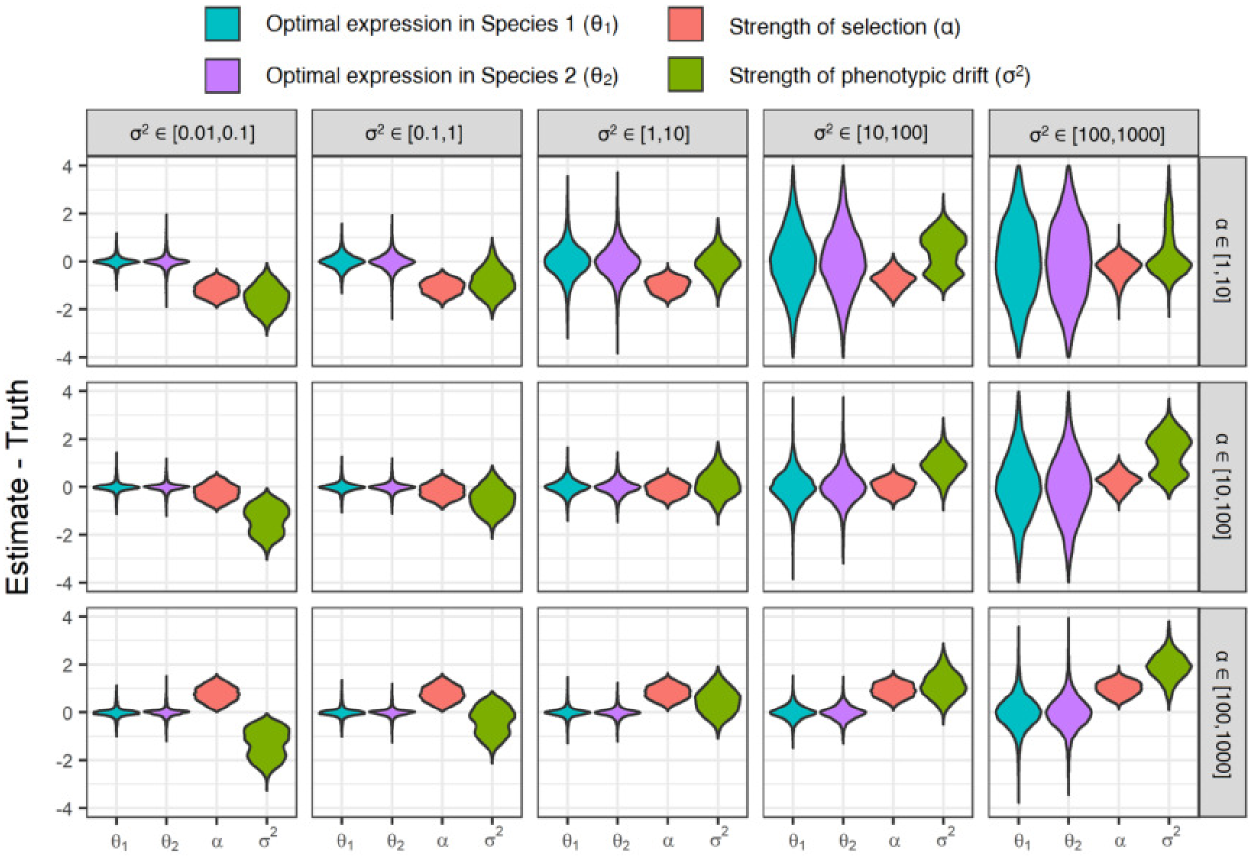


Figure S7: Parameter prediction performance of the neural network architecture of `PiXi` applied to test data simulated under specific parameter ranges for $\alpha$ and $\sigma^2$. Violin plots display distributions of mean parameter prediction errors across the $m = 6$ conditions for each simulated test dataset.

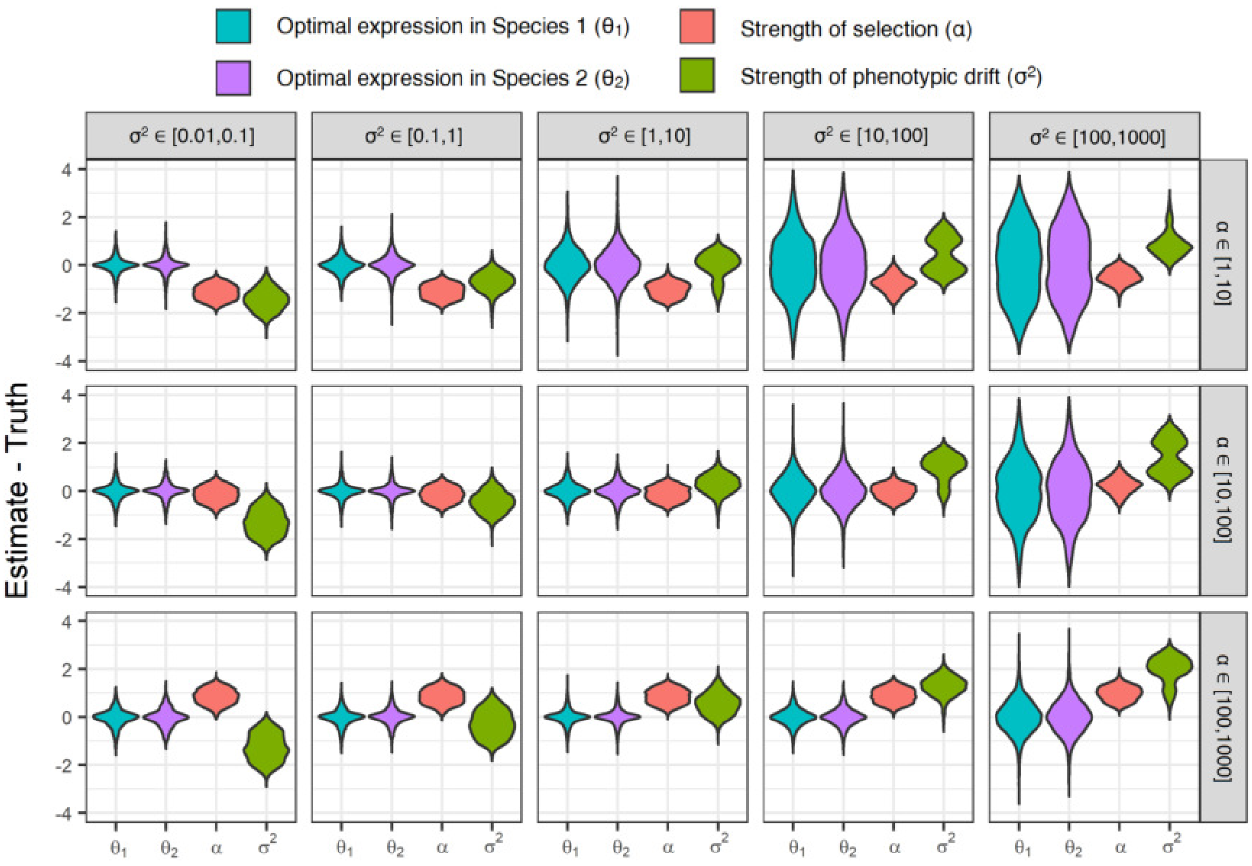


Figure S8: Parameter prediction performance of the random forest architecture of `PiXi` applied to test data simulated under specific parameter ranges for $\alpha$ and $\sigma^2$. Violin plots display distributions of mean parameter prediction errors across the $m = 6$ conditions for each simulated test dataset.

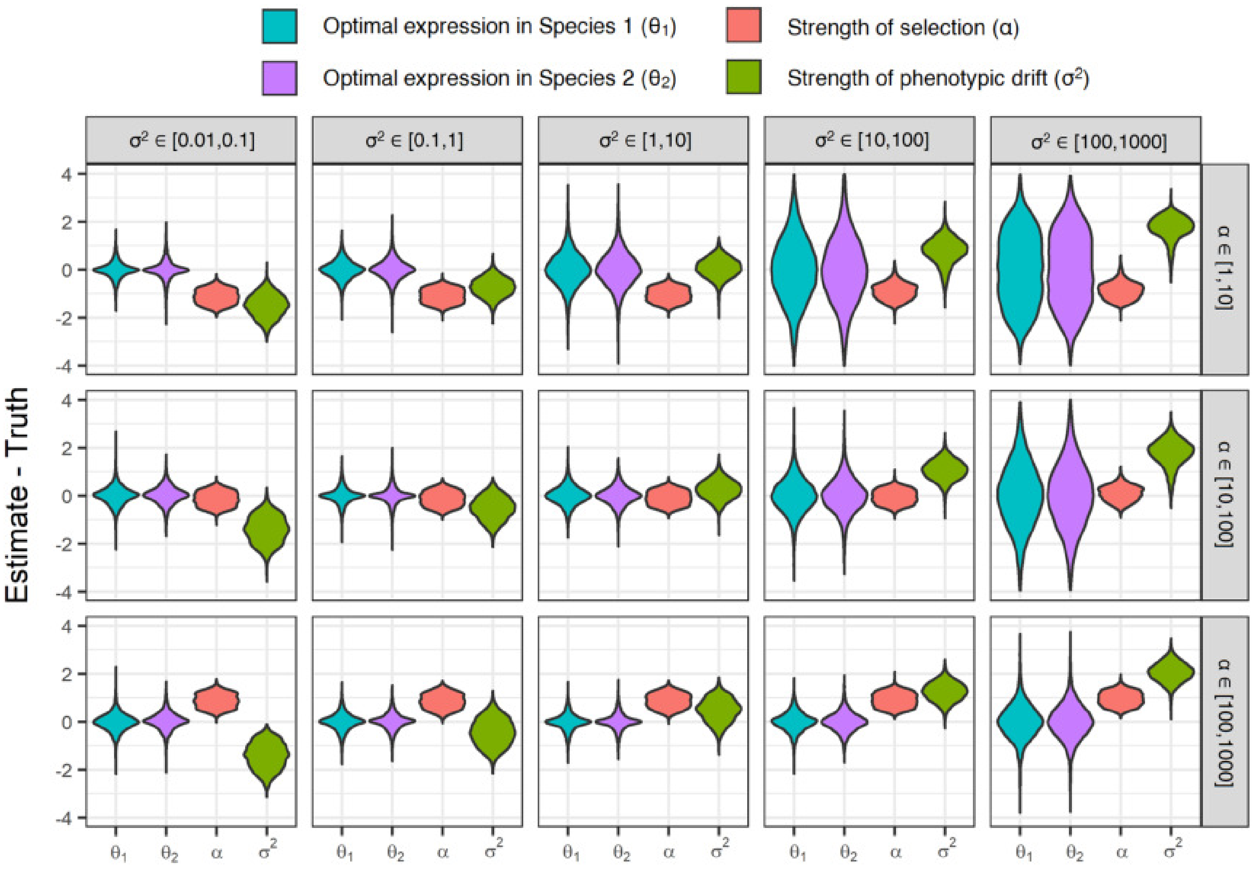


Figure S9: Parameter prediction performance of the support vector machine architecture of `PiXi` applied to test data simulated under specific parameter ranges for $\alpha$ and $\sigma^2$. Violin plots display distributions of mean parameter prediction errors across the $m = 6$ conditions for each simulated test dataset.

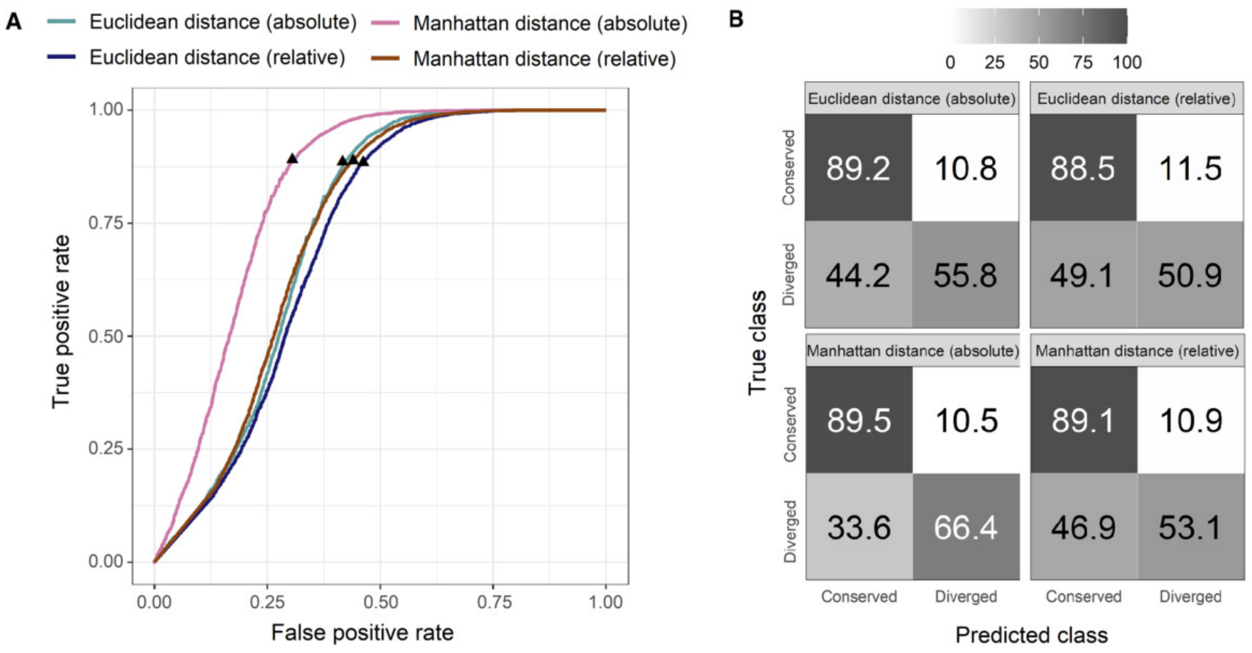


Figure S10: Classification performance of four distance-based classifiers applied to test data simulated under parameters $\alpha \in [1, 10^3]$ and $\sigma^2 \in [10^{-2}, 10^3]$ for each of the two classes. ($A$) Receiver operating characteristic curves showing the power of each method across the full range of false positive rates, with the black triangles depicting the cutoff chosen by cross-validation for the distance-based classifiers. ($B$) Confusion matrices depicting classification rates of the two classes for each method.
